# DAVI:Deep Learning Based Tool for Alignment and Single Nucleotide Variant identification

**DOI:** 10.1101/778647

**Authors:** Gaurav Gupta, Shubhi Saini

**Affiliations:** IIT Delhi

**Author notes:** Authors’ address: Gaurav Gupta,; Shubhi Saini,. **Reference Format:** Gaurav Gupta and Shubhi Saini. 2019. DAVI:Deep Learning Based Tool for Alignment and Single Nucleotide Variant identification (September 2019).

**Keywords:** genome, deep learning, single nucleotide variant caller, genome-assembly, pipeline, convolutional neural network, Recurrent Neural Network

## Abstract

The Next Generation Sequencing (NGS) technologies have provided affordable ways to generate errorful raw genetical data. To extract Variant Information from billions of NGS reads is still a daunting task which involves various hand-crafted and parameterized statistical tools. Here we propose a Deep Neural Networks (DNN) based alignment and SNV tool known as DAVI. DAVI consists of models for both global and local alignment and for Variant Calling. We have evaluated the performance of DAVI against existing state of the art tool-set and found that its accuracy and performance is comparable to existing tools used for benchmarking. We further demonstrate that while existing tools are based on data generated from a specific sequencing technology, the models proposed in DAVI are generic and can be used across different NGS technologies. Moreover, this approach is a migration from expert driven statistical models to generic, automated, self-learning models.

## 1 INTRODUCTION

Next generation sequencing [1] has opened up a new paradigm - It has provided affordable access to whole genome or genome sequences to many researchers which leads to personalized medicine and to personal genome project[2]. Many Mendelian disease studies have employed NGS to identify causal genes based on patient-specific variants[3]. Since a disease associated genetic variant rarely occurs among general healthy population, its interpretation in a patient is relatively simple. This interpretation simplicity by virtue of rarity of the variants however entails risk of false discoveries due to the errors in sequencing and false detection by variant calling methods. For success of clinical genomics and personalized medicine, fast and accurate identification of variant is most crucial. Thus NGS technology has shifted focus from generating genome data to swift and accurate information extraction. In NGS methods, a whole genome or targeted regions of the genome are randomly digested into small fragments (or short reads) that get sequenced and are then either aligned to a reference genome or assembled[4] to form complete DNA (de-novo assembly). From aligned reads, variants and indels are identified by segregating read errors from potential variants and then comparing contigs with reference genome. These identified variants are known as True Variants and the process of detecting variants in reads is known as Variant Calling. Since NGS represents a throughput technology, it is highly sensitive to technological errors and produces data which is erroneous, random, multiple and of high coverage. It is therefore required to clean and pre-process [5] [6] [7] [8] data before variant calling. Variants detection in a genome sequence is thus a multi-step process comprising of : (1) Mapping the reads to the indexed reference(alignment), (2) Sorting the reads based on their location, (3) Identifying and annotating the probable variant candidates, (4) Determining true variants and their genotypes from variant candidates with high confidence (variant calling). This makes the process of variant and indels (insertion and deletion of bases) identification highly dependent on reliable bio-informatics tools. There are many off-the shelf tools and pipelines available for variant identification[9]. These tools are based on string manipulation and statistical modeling and require parameter tweaking by domain experts. Moreover, different tool sets are required for different NGS technologies and for different species.

In this paper, we propose a novel deep neural network based tool “DAVI” (Deep Alignment and Variant Identification) which can be used across NGS technologies and species for raw reads alignment and SNV detection. We benchmark the computational performance of DAVI against state of the art GATK pipeline and demonstrate that it not only performs variant identification faster, but is also more specific than existing tool set.

## 2 CURRENT METHODOLOGY

As described above, all existing pipelines can be broadly divided into two major processes : One is aligning reads to reference **(Alignment)**, and the other is identification of variants from aligned reads **(Variant Calling)**.

Alignment is the process by which the NGS reads are aligned to their corresponding(most likely) locations in the reference genome. Alignment can be achieved by comparing reads with k-mers of reference genome. But due to imperfection in raw NGS reads(substitution and indels) an attempt to exactly match reads to reference k-mers will lead to rejection while comparing. Thus to perform alignment raw reads are divided into sub-reads and matching regions (contigs) with reference are identified. Matched sub-reads are expanded at contigs to find best suitable match of a read. There exist many tools which perform this local alignment like BWA, Novalign, Bowtie, BLAST etc. These algorithms use dynamic programming to locate region for best alignment. Algorithmically these algorithms can be divided into two main categories : (1) Hash Table based algorithms, indexing either the reads or the reference genome and (2) Burrows-Wheeler Transform based algorithms, using suffix trees and suffix arrays of the strings[10][4]. Recently there have been some attempts to use ML based algorithms for pseudo-alignment

Variant calling is the process by which variants are identified from the aligned sequence data with respect to reference sequence of the particular organism. GATK, VarScan2, Atlas-SNP2,SNVer are some of the tools that perform variant calling on aligned reads. SNP callers may also be divided into two different approaches: (1) Heuristic methods based on thresholds for coverage, base quality and variant allele frequency and (2) Probabilistic methods based on genotype likelihood calculations and Bayes’ theorem. Due to their computational demands, heuristic based methods are less commonly used than probabilistic methods [4].

### 2.1 GATK Best Practices Pipeline

For the purpose of benchmarking the performance of our proposed SNV identification methodology, we have experimented with Illumina2000 data. We have compared our results with the widely used *GATK best practices pipeline*[11] for SNV. GATK based pipeline has been selected because it gives best performance across existing variant calling pipelines against gold standard set of reference variant calls from GIAB[9]. This pipeline consists of BWA, Picard Tools and GATK packages.

GATK uses machine learning based algorithms in its *HaplotypeCaller* tool for variant detection, with *PairHMM* method for pairwaise aignment of each read against its haplotype. *PairHMM* is a pairwise alignment method that uses a Hidden Markov Model(HMM) and produces a likelihood score of observing the read, given the haplotype [12]. For each potentially variant site, the program applies Bayes’ rule, using the likelihoods of alleles given the read data to calculate the posterior likelihoods of each genotype per sample given the read data observed for that sample.

### 2.2 Deep Learning Based Approaches In Bio-informatics

Despite more than a decade of effort and thousands of dedicated researchers, the hand-crafted and parameterized statistical models used for variant calling still produce thousands of errors and missed variants in each genome [13]. Deep Neural Network based learning algorithms provide attractive solutions for many bio-informatics [14] problems because of their ability to scale for large dataset, and their effectiveness in identification of intrusive complex features from underlying data. There has been some success in area of de-novo peptide sequencing [15], mapping protein sequence to fold(deepSF)[16] and to predict protein binding sites in DNA and RNA(deepBind) [17]. CNN based models are used exhaustively in identification of Motif in DNA sequence [18], but not much has been done for the prediction of SNVs in raw DNA reads.

Only recently, Google has developed DeepVariant [19] tool which uses deep learning to predict variants in aligned and cleaned DNA reads. DeepVariant method uses CNN as universal approximator for identification of variants in NGS reads. It does so by finding candidate SNPs and indels in reads aligned to the reference genome with high-sensitivity but low specificity. The DeepVariant model uses Inception-v2 architecture to emit probabilities for each of the three diploid genotypes at a locus using a pileup image of the reference and read data around each candidate variant. There have been some attempts to use RNN for global alignment, but their results are not yet established [20].

In DAVI we have used both CNN and RNN algorithms for alignment. Using DNN we determine a DNA read comparator function (global alignment) or sub-read comparator function (local alignment),which determines best alignment based on scoring function. For Variant identification DAVI uses CNN like DeepVariant but instead of using pileup images with inception-v2 architecture, we use Position Specific Frequency Matrix(PSFM) to identify possible variant sites and a set of images(reference, substitution and indel) to predict variant and its genotype at a given location. The detailed model designs and experiments are illustrated in section below. We have used Illumina data of Ecoli (*CP012868.1 Escherichia coli str. K-12 substr. MG1655, complete genome*) and human data for validation of our experiments. We have also used pacBio human genome data for generalizing our model across NGS technologies.

## 3 DEEP ALIGNER AND VARIANT IDENTIFICATION

### 3.1 Alignment

As discussed above, to perform alignment we have to derive a *comparator function* which considers complete raw NGS reads (global alignment) or part of reads (local alignment), and reference k-mers from a location as input and performs comparison to output rank of similarity since perfect matches to reference sequence is elusive. Our *comparator function* should be able to handle some percentage of variation between two input streams while measuring degree of similarity. Higher similarity value implies better alignment at that location.

As we know, DNN works very well in extracting feature while being resilient over noise(variations). Moreover, since DNN based algorithm scales with data, our alignment use-case is a good fit for these algorithms. Two most widely used DNNs are Convolutional Neural Network(CNN) and Recurrent Neural Network (RNN). While CNN is generally used to extract features from local patches of data, RNN is used for sequential data where current outcome is dependent upon previous learnt patterns. We have explored both these techniques to solve our problem of alignment. The dataset used for training and testing of our DNN model is discussed in following subsection.

### 3.2 Dataset

To perform alignment we have used genome sequence of *Ecoli K-12*. For this reference genome we have used two sets of input reads: (1) Actual NGS reads, which generated from ILLUMINA sequencer and have length of 30-300bps with error rate of 1-2% and coverage of 30x. We called this dataset *real dataset*. We have used this dataset primarily for testing our models. (2) To learn a robust comparison function we have trained our models with what we call as *simulated dataset*. Reads for this dataset are generated from reference genome with max error rate of 40%. The process of generation of simulated reads and preprocessing of strings for alignment is explained in Appendix B.

### 3.3 DNN for Alignment

#### 3.3.1 Deep Conflation Based Model

In this model, we have used CNN to predict the most probable location of a given read by breaking the reads and reference into fragments of size k, called as k-mers. Let *r*_*i*_ be a k-mer obtained from reference location i. We calculate the similarity between k-mer of a given read w.r.t. several different reference k-mers from different locations < *r*_1_, *r*_2_, .., *r*_*i*_, …, *r*_*n*_ >. The highest similarity reference k-mer can then be extended in a fashion similar to BLAST algorithm. We formulate the problem of k-mer string matching as analogous to that of identifying match between a word and its misspelled variants. This formulation is similar to conflation problem of business data.”Character-level Deep Conflation For Business Data Analytic” [21] discusses the model developed to solve the conflation problem. We have extended this model for our application. The details of model used for Alignment are discussed in Appendix B. Figure 2 Gives an overview of the model.

**Fig. 1.**
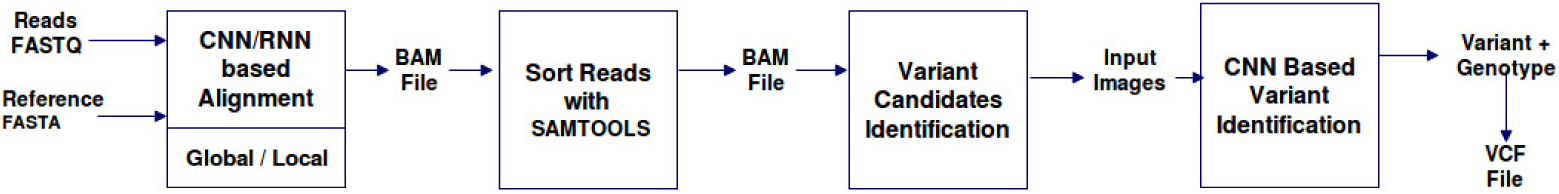
DAVI pipeline.

**Fig. 2.**
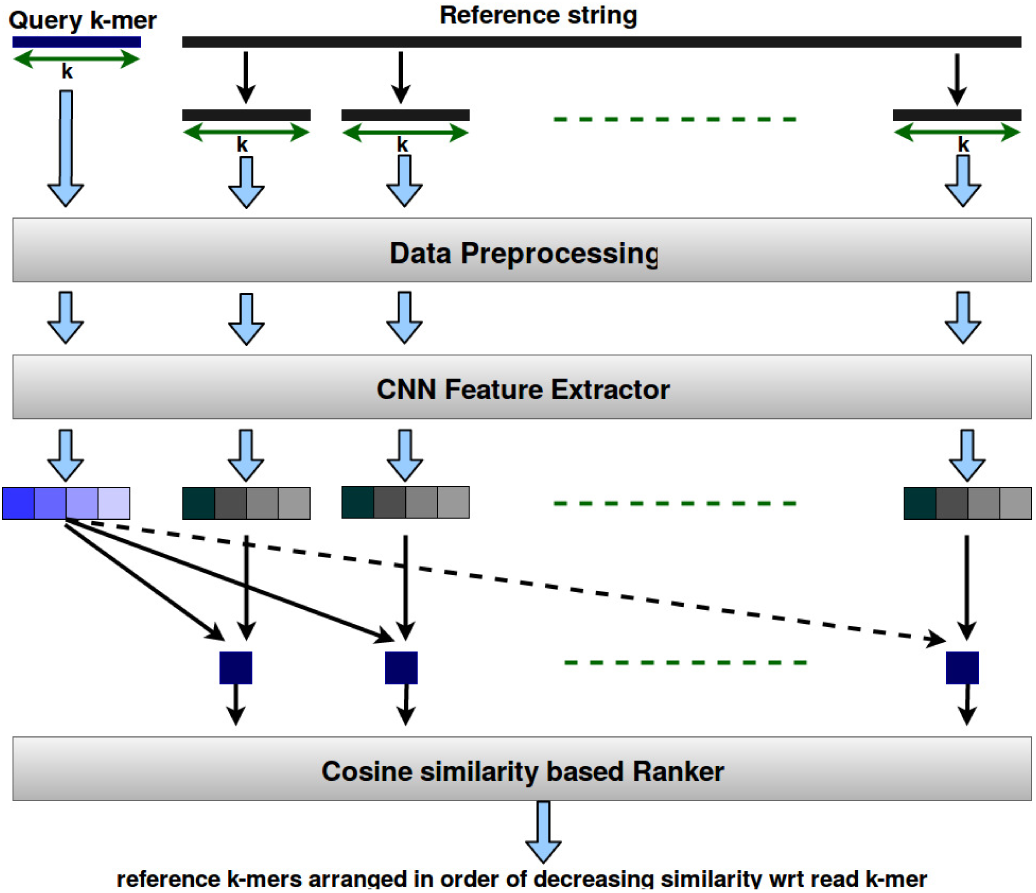
Deep Conflation Based Model.

##### Evaluation on Simulated Dataset

To perform alignment using this model, a simulated dataset was generated from genome sequence of ecoli(*CP012868.1 Escherichia coli str. K-12 substr. MG1655, complete genome*). We experimented by generating a set of 10000 random 100-mers containing characters A,C,G and T. For each 100-mer, we also generated a 100-mer with 40% Insertion,deletion and substitution starting at a random location. After training the model for 3 epochs on entire dataset, with mini-batches of size 100 the model was able to achieve a testing accuracy of 99.4%.

The entire dataset was divided into training and testing data, using a split ratio of 90:10 for training:testing. The training data was further split in ratio of 90:10 for training and validation, with data selected through random permutation on dataset for each epoch. With this splitting, each epoch contains 81 minibatches. Figure 3 shows the change in Loss and accuracy of prediction over number of minibatches during training. Accuracy for validation and testing is measured as the number of mutated strings for which the correct reference was ranked highest in the given set.

**Fig. 3.**
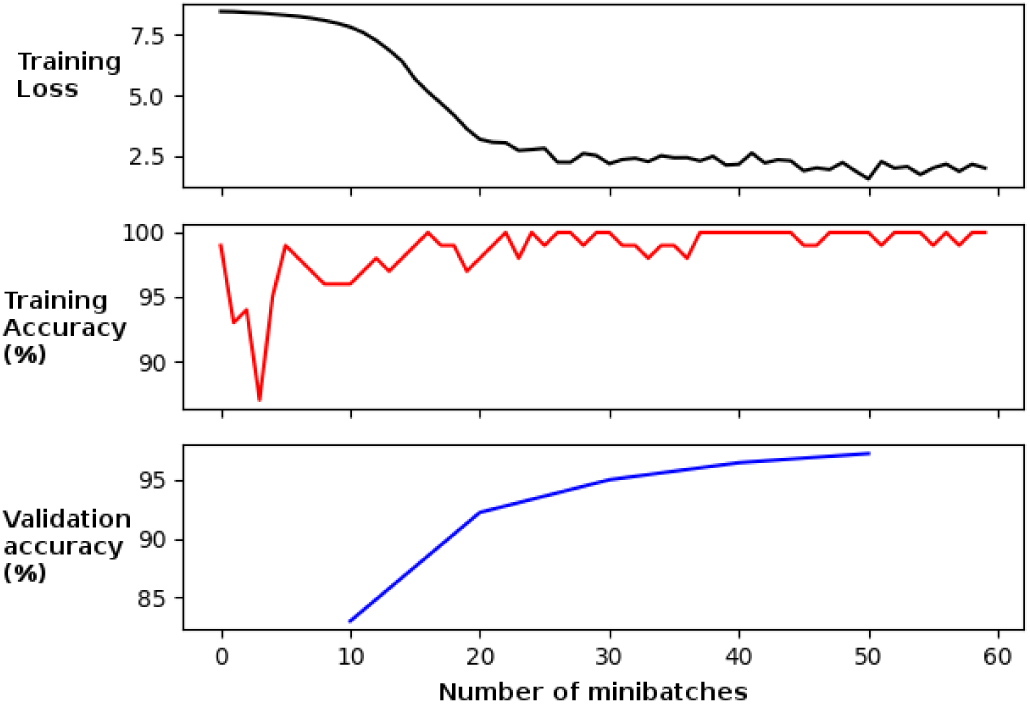
Loss vs number of minibatches.

##### Evaluation on Real Data

To test the accuracy of network trained with above simulated dataset with real data, the following procedure was used. We have taken EColi reference same as that used in simulated data and NGS reads from *SRR2724094_1.bam* (fastq aligned by BWA-MEM). The process is defined in Algorithm ?? (Appendix A). Model was tested on 100 batches of the above dataset where input sequence length is of 100 nucleotide bases. Statistics of test are shown in Table 3.3.1 and Fig.3.3.1.

##### Observations

Deep Conflation model has been found to be useful for predicting the best match reference k-mer to a given read. In this model the length of read (k-mer) does not affect the accuracy of prediction. Thus this model can be used for global alignment, since a given read of length ‘r’ can be aligned by searching for best reference k-mer of same length by straightforward comparison. Since feature extraction for reference k-mers is required only once, we create a database of reference features first time. Subsequently, to test the alignment of reads against this reference, the features are queried from the database.

### 3.4 RNN Based Alignment

Recurrent Neural Networks are memory based Neural Networks. They are especially useful with sequential data because each neuron/unit can use its internal memory to maintain information about the previous input. The two most widely used RNNs are : Vanilla RNN and LSTM (Long Short Term Memory). RNN is estimated to be faster network as compared to LSTM, but it falls short in memorizing and representing long term patterns associated with input data stream. Thus, to determine which Recurrent Neural Network will be feasible for comparing genome strings, we modeled both RNN cells and LSTM cells. We further improved these models using *CoDeepNEAT* techniques.

#### 3.4.1 Architecture

Alignment of query reads with respect to a given reference sequence, works by comparing query string with a series of reference strings, and determining the degree of confidence of their alignment. Since a read may be aligned to more than one locations in the reference, ‘degree of confidence’ gives a measure to choose the best site for alignment. To perform alignment using neural network, our RNNs takes reference string and query string as input and then classifies them into one of the two classes: namely, *Matched* class and *Unmatched* class. Then from all Matched cases based on best matching score, a reference string is picked for alignment. For classification, we have modeled our DNN as shown in Fig 5. The value of hyper-parameters used in Vanilla RNN and LSTM model are listed in Table 2.

**Table 1.**
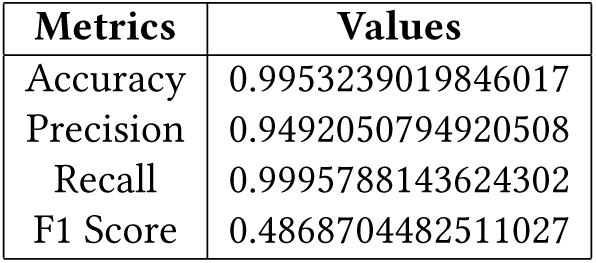
Performance Metrics of Deep Conflation Based Alignment Model.

**Table 2.**
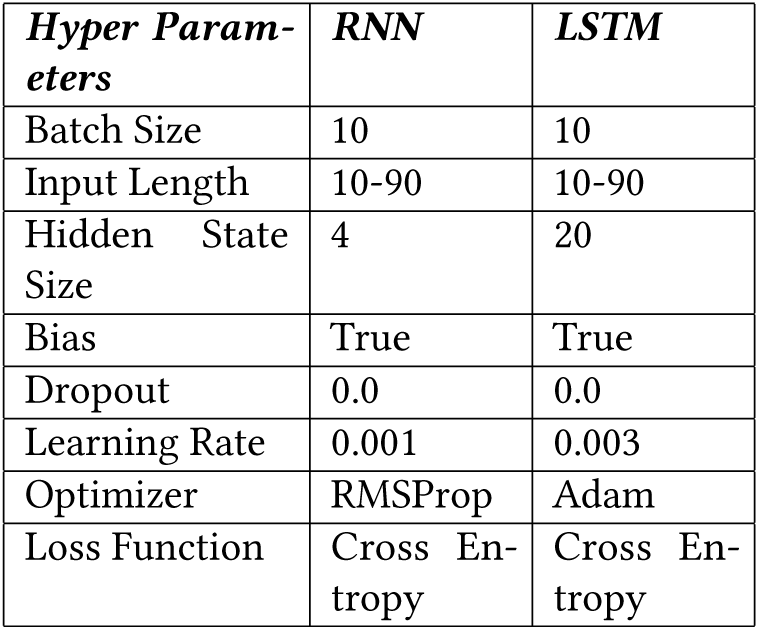
Vanilla RNN and LSTM Hyper Parameters.

**Fig. 4.**
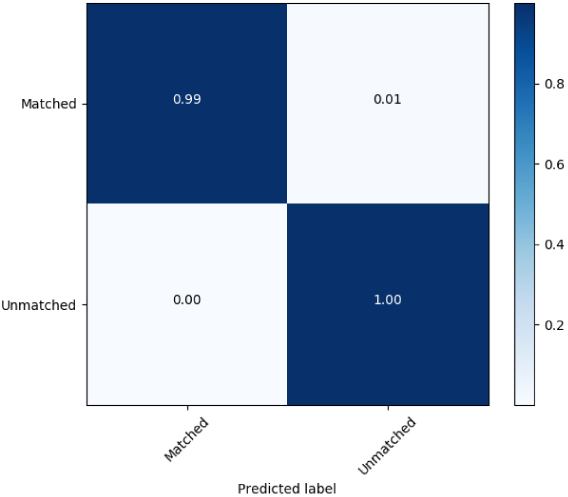
Confusion Matrix for Deep Conflation Alignment Model.

**Fig. 5.**
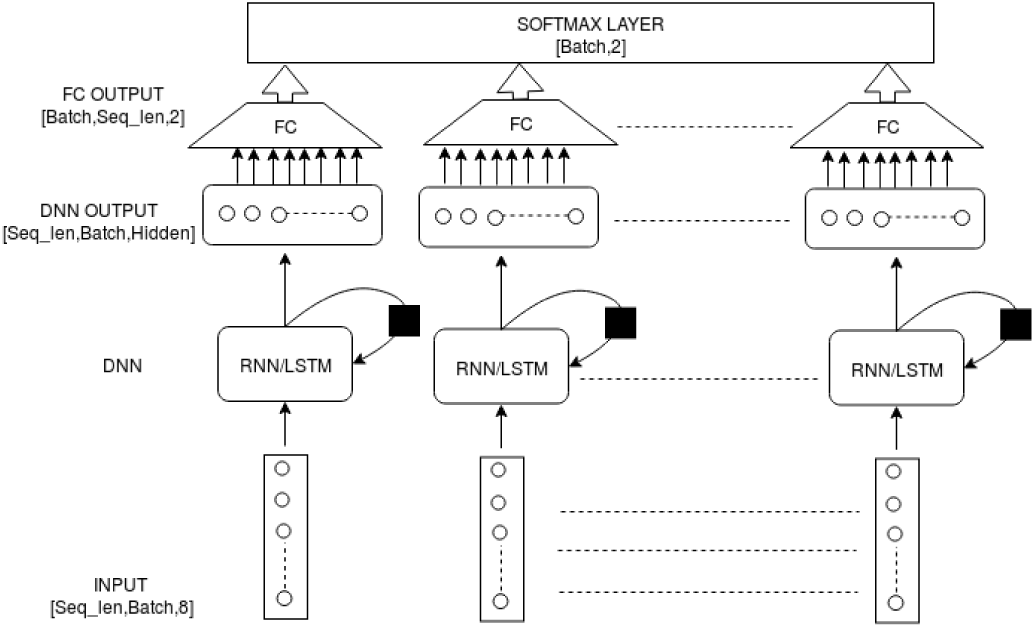
RNN Based Alignment Model Architecture.

#### 3.4.2 Experiment

Models of both RNNs are trained on Simulated Dataset. Input to model is generated by concatenating raw sequence of fixed length with reference sequence of same length against which comparison to be made. This concatenated sequence is then converted to Hot vector Encoding to generate input for models. For Vanilla RNN 50 epoch with 380 batches of training dataset is used while for LSTM 100 epochs are used for training same dataset. To observe over-fitting, validation test was performed on validation dataset at regular interval.

##### Evaluation and Observations

Trained RNN models was evaluated on test dataset of simulated dataset. Mean accuracy of 87.55% was observed for Vanilla RNN, The accuracy graph and model test parameters of Vanilla RNN for test dataset is shown in Fig. 6. While mean accuracy of 79.6 % is observed for LSTM. The accuracy graph and model test parameters of LSTM for test dataset is shown in Fig.7. Detail Design for both Vanilla RNN and LSTM with training accuracy is illustrated in Appendix C

**Fig. 6.**
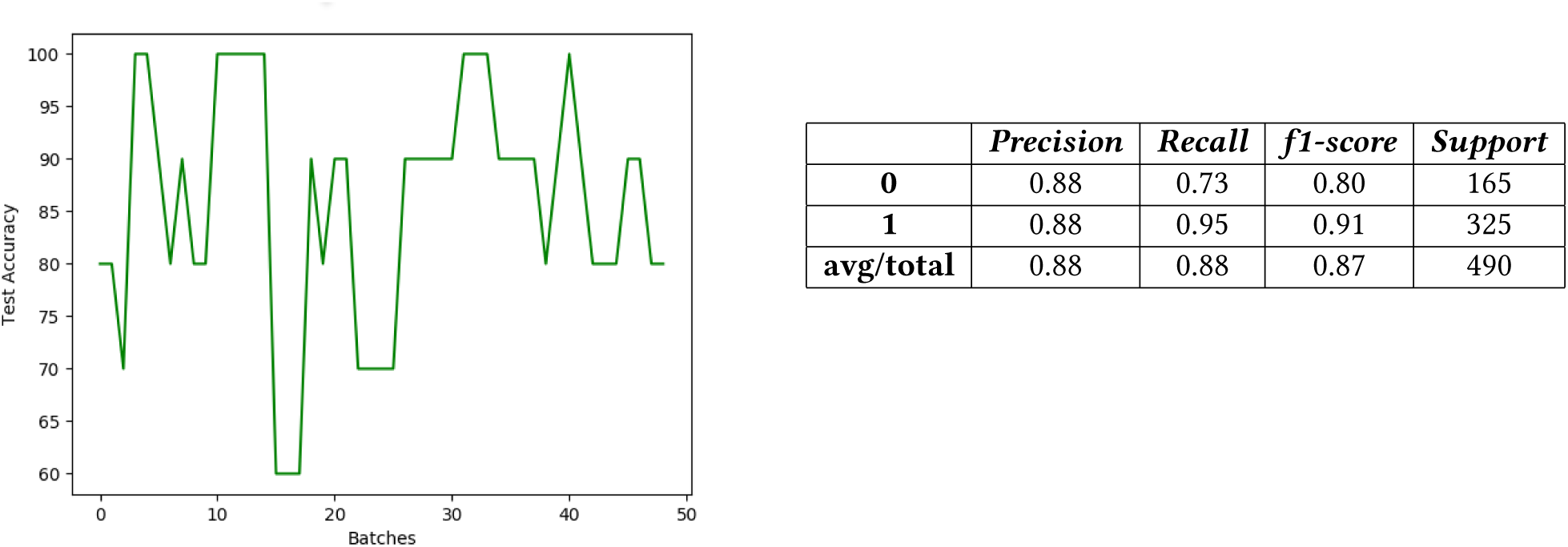
RNN Model Test Accuracy.

**Fig. 7.**
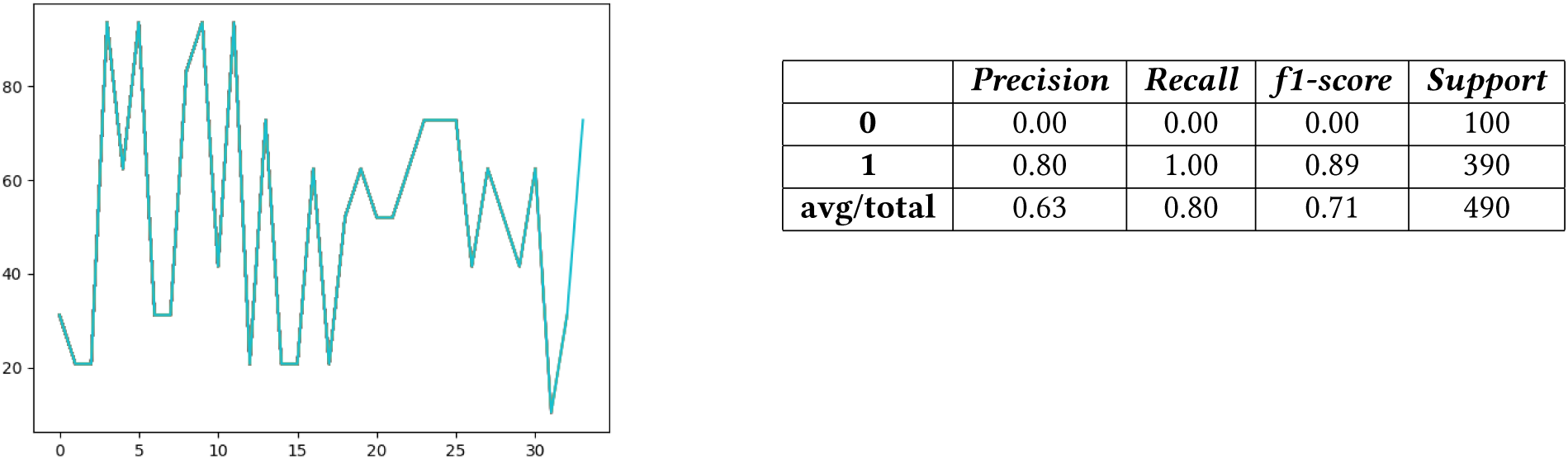
LSTM Model Test Accuracy.

##### Comparison of RNN Models

We have performed experiments with different sequence lengths ranging from 10 to 90. The number of hidden units in models is kept at 1/3 of sequence length. We have observed that as size of sequence grows beyond a certain threshold, both RNN models show no learning and accuracy drops significantly. For Vanilla RNN, model threshold reaches for sequence length above 50, while for LSTM model threshold is reached at length of 80. The test accuracy of Vanilla RNN model is significantly higher than LSTM model and also Vanilla RNN model trains faster as compared to LSTM model. Test accuracy of both models vs sequence length is shown in Fig. 8.

**Fig. 8.**
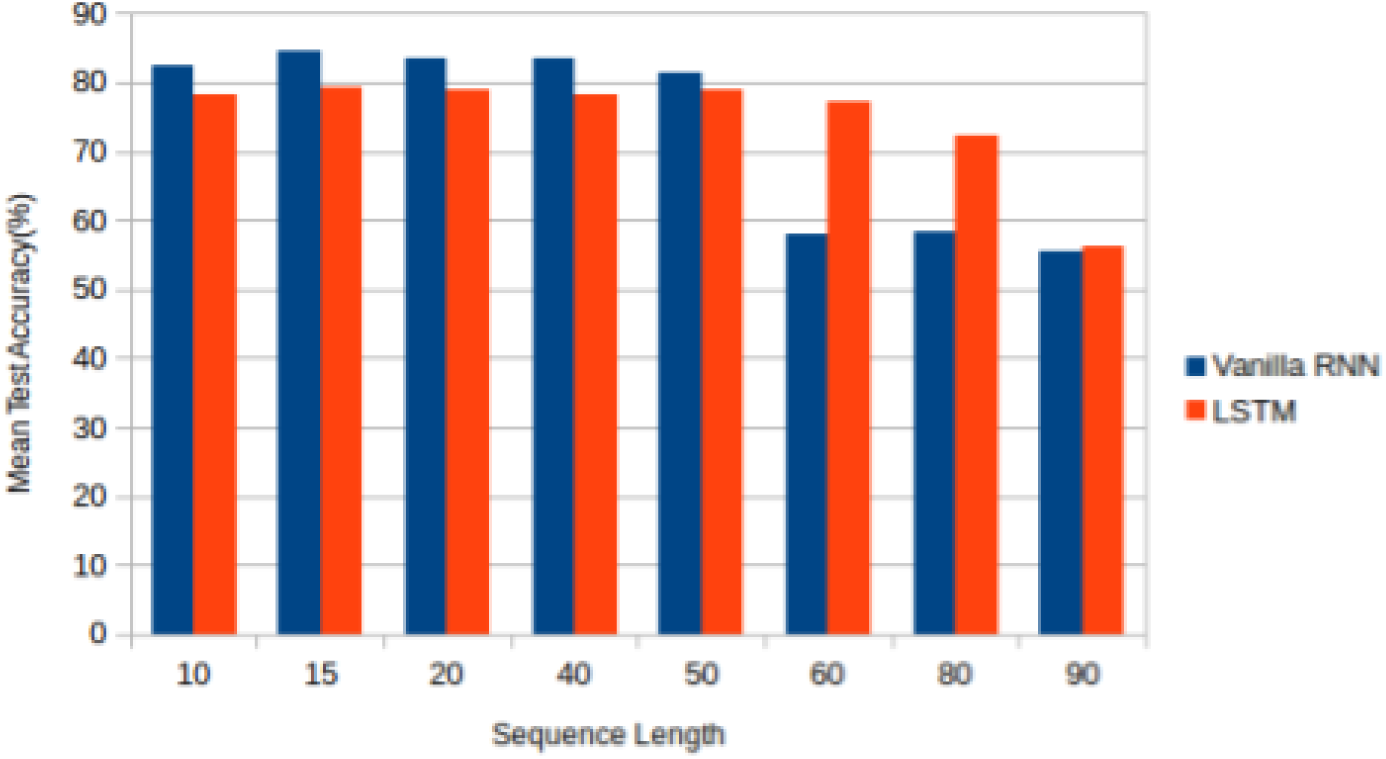
Test Accuracy Comparison of Vanilla RNN and LSTM model with Input Sequence Length.

As observed Vanilla RNN model has better accuracy as compared to LSTM for query length up to 40bps, we have used Vanilla RNN for alignment. Since query length of NGS data is more than 40bps, Vanilla RNN based model can be used for *Local Alignment* only.

### 3.5 Optimization of RNN models

The above two experiments are clear indicators that vanilla RNN based Neural Network does perform well for genome comparison, but designing an optimal RNN network is a difficult task as each network has large number of hyper parameters to optimize. To automate the process of finding best values for each hyper-parameter we have used Neuro Evolution technique known as CoDeepNeat[22]. CoDeepNeat uses existing neuro evolution technique of NEAT [23], which has been successful in evolving topologies and weights of relatively small recurrent networks. The fitness of evolved DNN is determined by how well system is getting trained by gradient descent, to perform its task. CoDeepNeat uses genetic algorithm for optimization which is combined with gradient descent to evolve large and complex DNN.

#### 3.5.1 Extending CoDeepNeat for RNN

In our implementation of CoDeepNeat, population of chromosomes is created by randomly initialized hyper parameters of network and slowly evolving them though mutation. Instead of representing each chromosome as a layer in DNN, in our design each chromosome itself is represented as a DNN. Advantage of this modification is that it nullifies the requirement of crossover in genetic algorithm as crossed over offspring can be generated by the application of mutation only. Although this approach limits the size of DNN and number of different layers that can be stacked, yet this is not a constraint since our Network is small. To evolve the networks, we have used two different mutation operators, namely Random Mutation Operator and Gaussian Mutation Operator [24] To evolve population offspring with selection, we have used DNN learning as fitness function. In particular, accuracy of validation test of DNN after a fixed number of epochs. Chromosomal population is evolved using genetic algorithm. The algorithm used for evolving chromosome population selects three elites from parent generation and are evolved in next generation without mutation while all other off springs are mutated version of previous generation chromosomes. The detailed algorithm for evolution is described in Algorithm 4 (Appendix A).

#### 3.5.2 Evolution & Training

Evolution was carried out for 10 generations(epochs) and best DNN was selected as the one with highest accuracy in any generation of evolution. To mutate chromosomes, mutation operators were applied to each hyper parameter with probability of 0.2. Since training a DNN is computationally expensive and size of population was 30, each network was trained for only two epochs on the training set. Validation Set was used for calculating fitness of DNN. Due to small number of epochs validation accuracy was calculated once, at the end of training. It is argued that due to small number of epochs, the network which is evolved is the one which trains fastest rather than the one with highest accuracy. Thus, through this technique we have automated the design of DNN topologies which gives model with shortest training time with relatively higher accuracy.

#### 3.5.3 Experimental Setup and Evaluation

System is built using *Pytorch* libraries with *Tensorflow* backend for CUDA platform on single GPU. Time taken for model to evolve is 49.32 hrs. After network is evolved for 10 generations, for each DNN (chromosome) in population, mean validation test accuracy is calculated. Fig. 9 shows the performance of population across generations. It is clear from this figure that system evolves till 6th generation by increasing mean accuracy and minimizing the standard deviation of population. But after 6th generation, mean accuracy decreases due to random mutation and standard deviation of population increases. During evolution most of the DNNs (population) trained quickly and had accuracy between 90%-100% as shown by density of clusters in Fig. 9. For the optimal solution, we chose DNN from 6th generation which provides maximum accuracy. DNN with highest accuracy and smallest standard deviation has been picked up as best DNN for nucleotide base comparison. The values of hyper parameters of best DNN suggested by model are listed in Table 3.

**Table 3.** Hyper Parameter Values of Suggested RNN.

| *Hyper Parameter* | *Values* |
| --- | --- |
| Number of Layers | 1 |
| Learning Rate | 0.0938099 |
| Optimizer | RMSProp |
| Bias | False |
| Hidden Size | 7 |
| Dropout Rate | 0.2653750429101209 |
| Activation Function | sigmoid |
| Bi-directional | False |

**Fig. 9.**
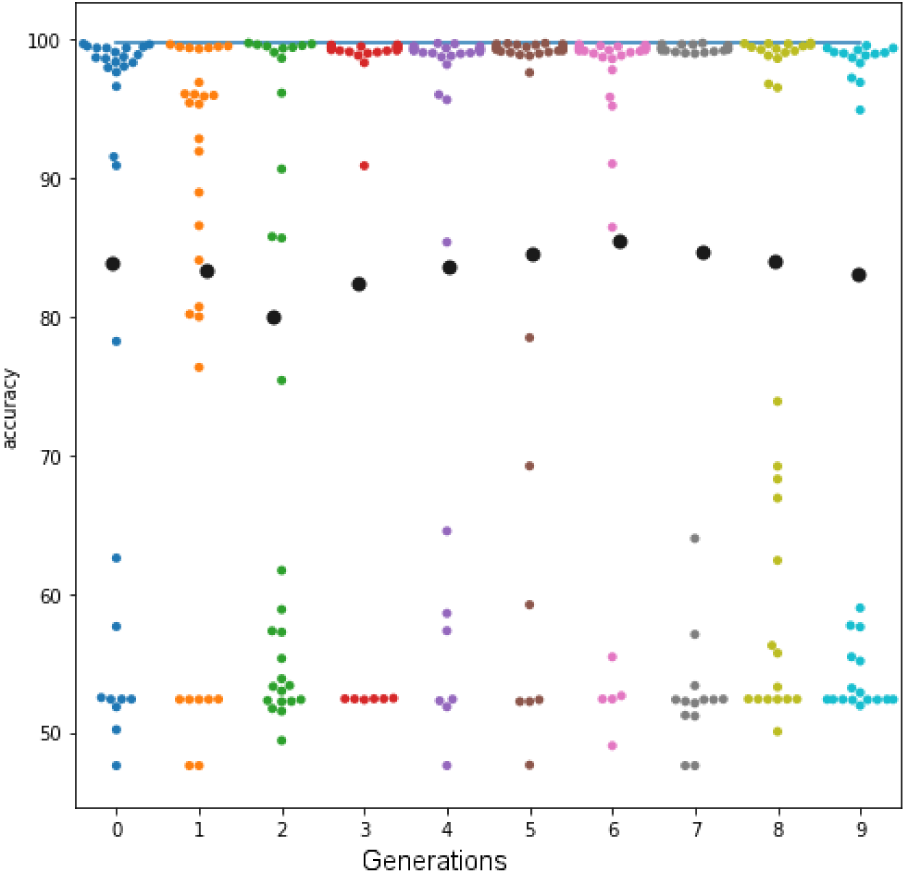
Accuracy of population after each Generation.

#### 3.5.4 Evaluation Of Optimized RNN with Simulated Data

To train and test optimized RNN model, simulated dataset is used. Training Loss graph (Fig. 10) of optimal DNN converges faster as compared to Vanilla RNN. Vanilla RNN learning starts around 3500 batches while in optimal DNN learning starts just after 2500. Hence, number to batches required for training optimal DNN is much less than Vanilla RNN model. Mean test accuracy of optimized RNN for 100 samples of simulated dataset is 96.123 % as compared to 84.55 % of Vanilla RNN (Fig.11). When tested on real dataset with input sequence of length 15 nucleotide bases mean accuracy for 100 samples increases to 98.88. The accuracy plot and report are shown in Fig 13 and 12

**Fig. 10.**
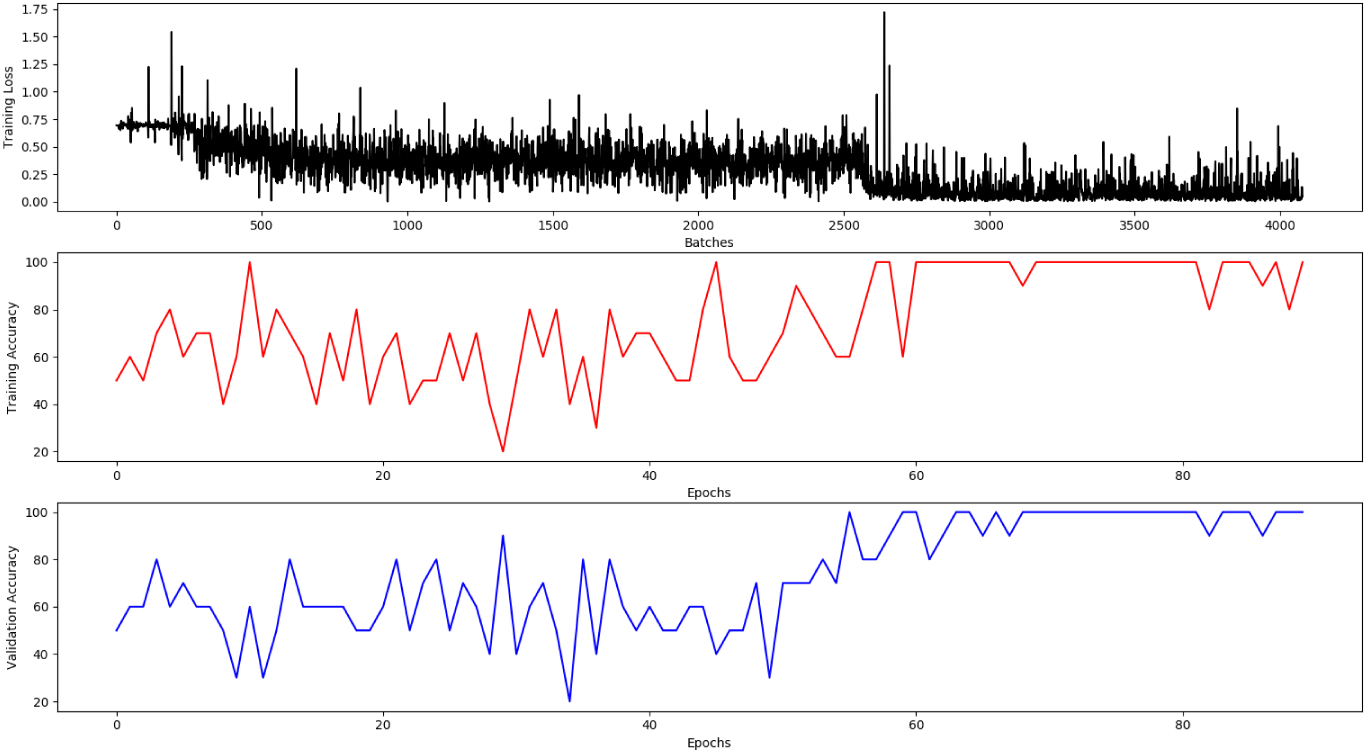
Performance of optimized RNN.

**Fig. 11.**
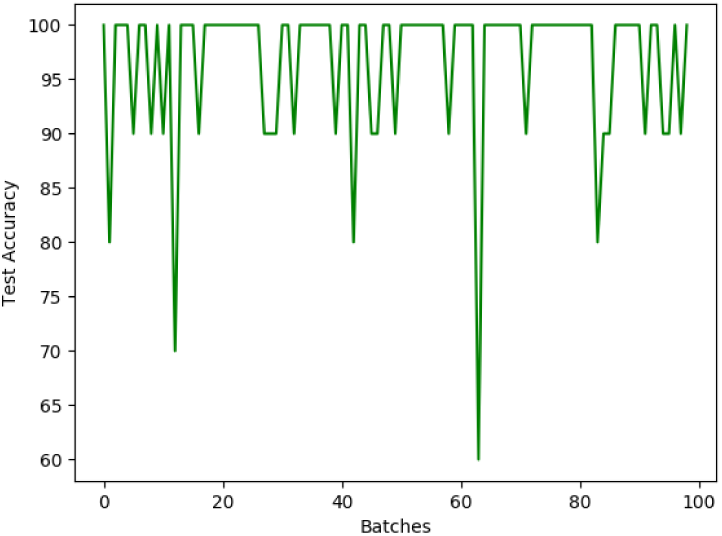
Test Accuracy of Optimized RNN.

**Fig. 12.**
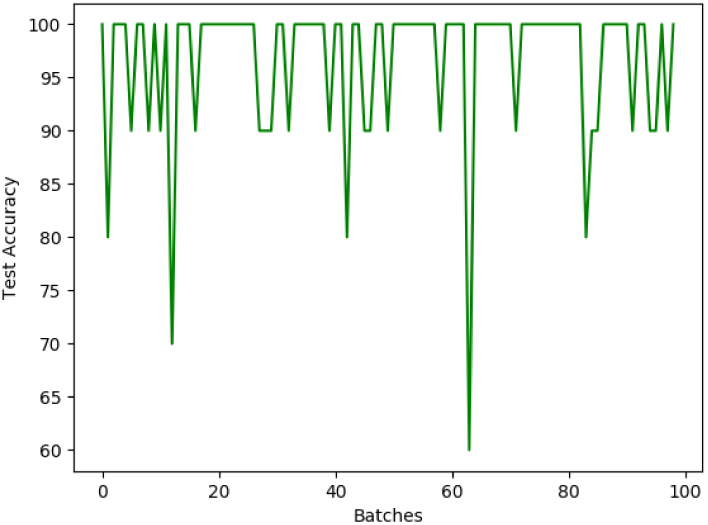
Accuracy Of Optimized RNN on Real Dataset.

**Fig. 13.**
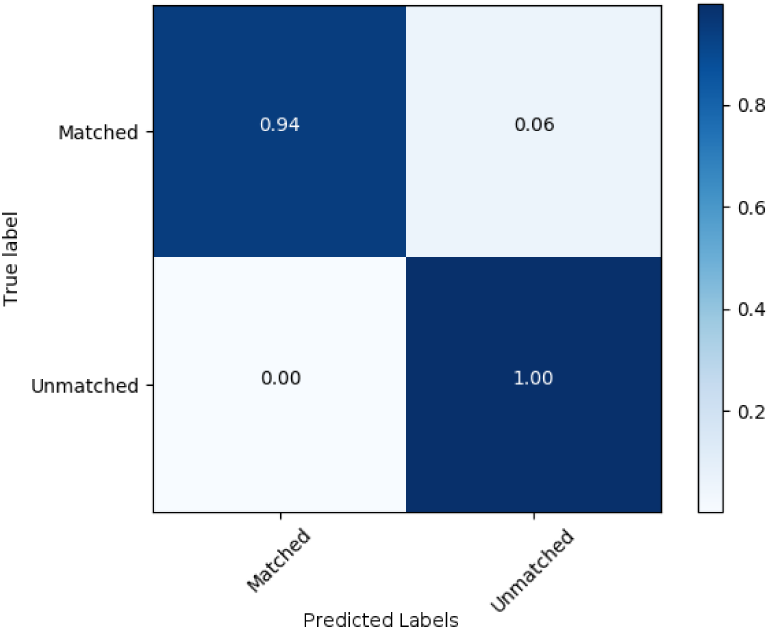
Confusion Matrix Of Optimized RNN On Real Dataset.

##### Limitation and Assumptions

Limitations of RNN based model are as follows:

- We have assumed the reads to be of fixed length and maximum length of a read is lesser than 40 bps.
- This model falls short in doing global alignment as, length of read generated by Illumina NGS are greater than 40 bps.
- We have not considered quality of reads while performing comparison.
- We have assumed maximum errors in high quality reads are 40% of total length.

## 4 USING DNN MODELS FOR ALIGNMENT

We have described CNN and RNN based models which can be used to generate *comparator function* for alignment. These models need to be discussed from the viewpoint of their usage in Alignment framework. RNN based alignment model, on the other hand has been found to work well on k-mers with length less than 40bps. Since the reads produced by Illumina are greater than 40bp in length, this model is more suited for local alignment of reads. To align a read w.r.t. reference genome, RNN has to be used in consultation with Local Alignment Algorithm as described in Algorithm 5 (Appendix A).

To increase the speed of alignment, we have used heuristics similar to BLAST to reduce our search space, by reducing the number of comparisons. Instead of comparing with each reference k-mer, the algorithm uses 5-mers as seed, and based on the seed all possible locations are searched for alignment. To reduce search space for query sequence, a database of all possible 5-mers with their locations in reference genome are stored in dictionary format. Once database is created, all possible 5-mers in query sequence are read. These query 5-mers are then searched in database and their corresponding location of occurrences are saved as probable locations for alignment. The detailed work flow of algorithm is as described in Algorithm 1.

### 4.1 Comparison of Models

We have experimented with Ecoli data on Deep Conflation and RNN models as described in Algorithms 3, 5 and 1. EColi Reference used was the same as that across all alignment models, and reads were obtained from sample *SRR272401_1*. The models were compared for time taken by them to train and predict accuracy, and results are summarized in Table 5.

**Table 4.** Performance Metrics of Optimized RNN Alignment Model.

| *Metrics* | *Values* |
| --- | --- |
| Accuracy | 0.9883333333333333 |
| Precision | 0.9902534113060428 |
| Recall | 0.996078431372549 |
| F1 Score | 0.4965 |

**Table 5.** Alignment Models Timing Comparison.

| *Model* | *Training Time* | *Prediction Accuracy* | *Alignment Type* |
| --- | --- | --- | --- |
| Deep Conflation Model | 4 hours | 99.5 | Global |
| RNN Model | 14.24 min | 98.8 | Local |

The models proposed for alignment are also compared with existing state-of art alignment tools, mainly with BLAST and BWA-MEM. For BLAST [25] we have taken open source python implementation while for BWA-MEM standard tool was taken for comparison. The result of performance comparison is shown in Table 5. It is observed that BWA-MEM is highly optimized for alignment and it is fastest irrespective of query length. BLAST is also optimized by using several heuristics to limit the search space. Our proposed model Recurrent Aligner works well for query of small length as model is designed for optimum length of 32bps. While Deep Conflation Model works well for query length based on training configuration. When trained for 100bps sequences, the model works well for sequences 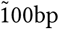 length, but is unable to align queries shorter than 50 bps. On training the model with 30bps sequences, queries of 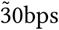 are aligned correctly. Therefore deep conflation model is not able to align queries with more than 50% of padding. For high quality alignment, threshold value of score can be set in proposed model.

#### Algorithm 1 Alignment with Heuristic

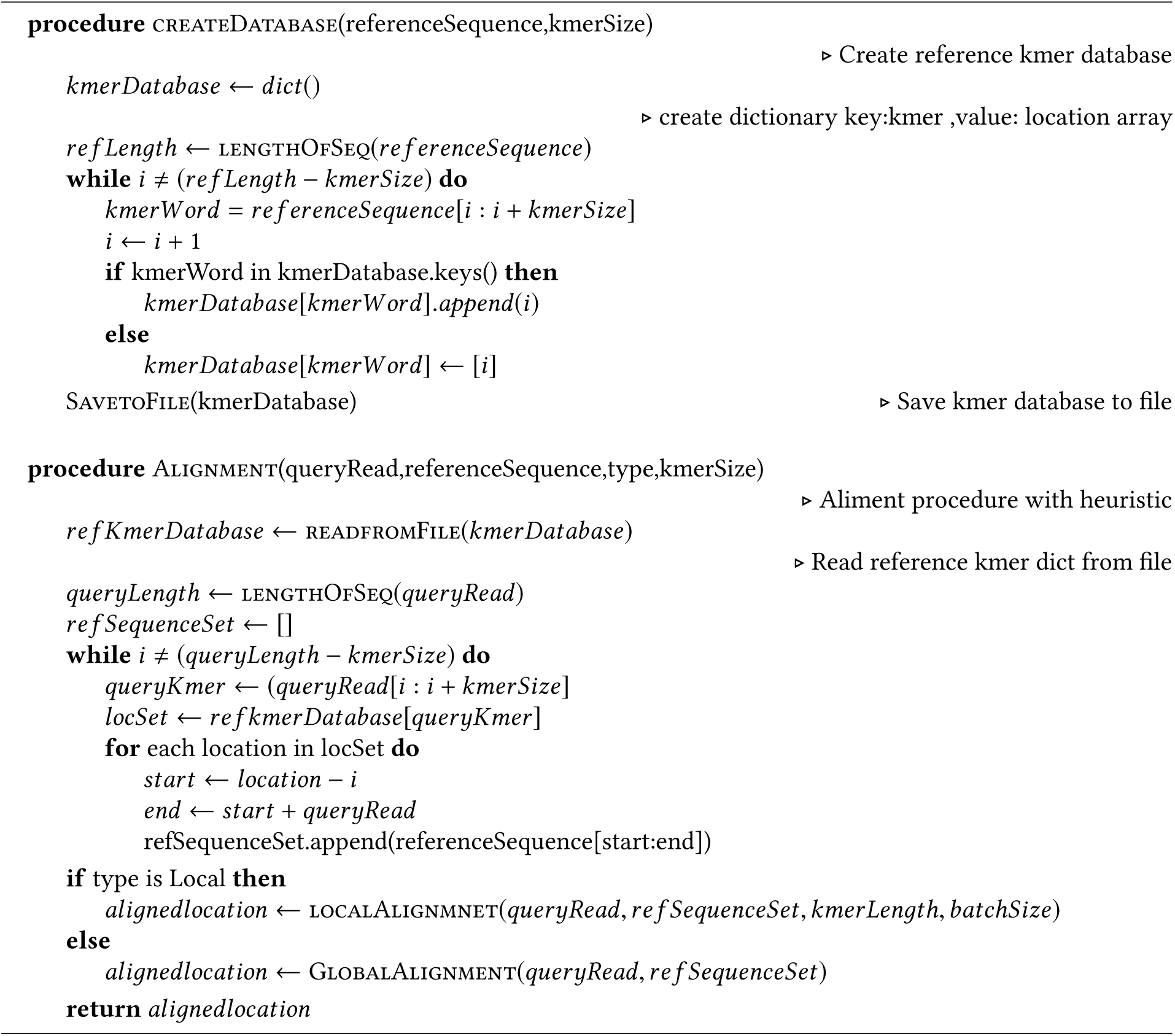

### 4.2 Deep Learning Based Variant Calling

The SNV and indel identification in a genome is a complex problem, which can be broadly broken down into two subproblems: Identification of variant and Classification of their type. Probable Variant sites can be classified into four categories : Heterozygous, Homozygous, Non-variant and Non-SNP. Since we have already discussed how CNN based deep learning model can be used for extracting feature and for classification of genomic data, we have used CNN for identification and classification of variants. The proposed Variant Caller pipeline is as described in Fig.14. Data is preprocessed where from aligned and sorted reads probable variant candidates are located using Position Frequency Matrix. These variant candidates are converted to Input Image Matrix, which is fed to Trained CNN for Classification of Probable Variant Candidates to Variant/Non-Variant and their genotype. The details of data processing, input image creation and label data generations is described in Appendix D For training the CNN model, we have used two sets of datasets, one of human genome and another of Ecoli. For human, we have used Genome in a Bottle reference dataset [26]. Out of this dataset we have used *NA12878* [27] data, in which input data is in BAM format while variant data is in VCF format. To consider variant which has high confident, interval data was used in BED format.

**Fig. 14.**
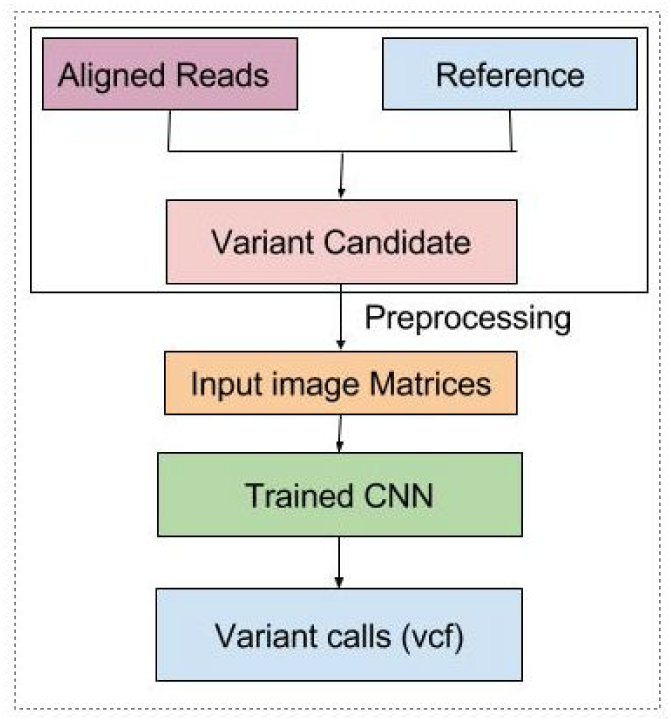
Proposed CNN based Variant Caller.

#### 4.2.1 Convolutional Neural Network

The network consists of two convolution layers with max pool layers and three densely connected hidden layers. In convolution layer, kernels of size (2,4) and of size (3,4) are applied to generate 48 convolution filters. Three densely connected layers use exponential linear activation function [28] with dropout rate of 0.05. The above network is trained using Adam Optimizer with learning rate 0.0005. The output layer contains two groups of output - For the first 4 output units, we learn about the possible bases of the site of interests. For example, if the data indicates the site has a base “C”, we like to train the network to output [0, 1, 0, 0]. If a site has heterozygous variants, for example,”A/G”, then we would like to output [0.5, 0, 0.5, 0]. We use mean square loss for these 4 units. For 2^nd^ group of outputs, the units consist of variant type. We use a vector of 4 elements to encode all possible scenarios. A variant call can be of any category as mentioned above. We use a soft max layer and use cross-entropy for the loss function for these 4 units.The architecture of CNN is depicted in Fig.15.

**Fig. 15.**
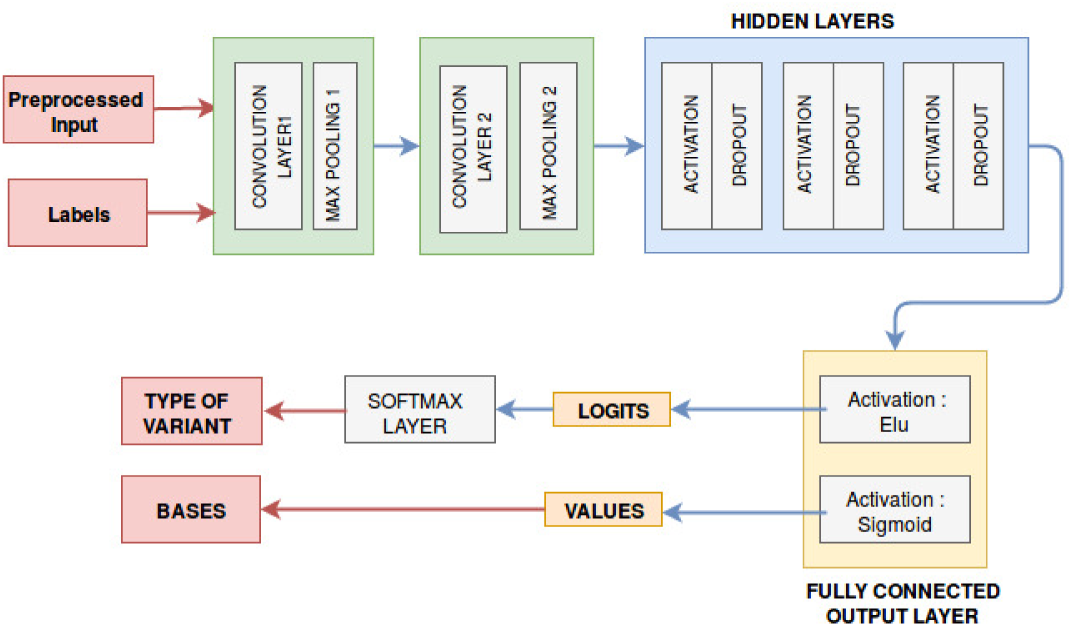
CNN architecture for Variant Calling.

##### Training

CNN was trained on data as mentioned above. Training images were sent as a batch of size 500, each image of dimensions 15* 4* 3. The first 30000 images were used for training and next 30000 were used for validation. Number of epochs for training was 3000. Time taken to train network on 30000 reads of *NA12878-chr21* was 284.289 seconds.

#### 4.2.2 Evaluation

To evaluate performance of our model we have generated confusion matrix for Genotypes of variants and report of our prediction as shown in Fig 16 and Table 8. It is clear from prediction results that our model output has high specificity and moderate sensitivity.

**Table 6.** Time taken (in sec) by DNN Models to Predict Location of Read.

| Read Name | Length of Read | RNN | Deep Conflation | BLAST | BWA-MEM |
| --- | --- | --- | --- | --- | --- |
| SRR2724094.1.1 1 | 300 | 4.34 | 1.8 | 10.85 | 0.64 |
| SRR2724094.3.1 3 | 37 | 0.624843 | 0.15 | 0.169 | 0.045 |
| SRR2724094.7.1 7 | 86 | 0.377 | 0.284 | 0.876 | 0.047 |
| SRR2724094.10.1 10 | 168 | 1.548 | 0.603 | 0.72 | 0.055 |

**Table 7.** Reads Location Prediction by different Models.

| Read | BLAST |  | RNN |  | Deep Conflation |  |
| --- | --- | --- | --- | --- | --- | --- |
|  | Location | Score | Location | Score | Location | Score |
| SRR2724094.1.1 1 | - | - | 4258852 | 0.388 | 4258852 | 0.389 |
| SRR2724094.3.1 3 | 3073798 | 0.945 | 3073797 | 0.95 | 3073797 | 0.2 |
| SRR2724094.7.1 7 | 306992 | 0.97 | 306991 | 0.98 | 306991 | 0.98 |
| SRR2724094.10.1 10 | 1564588 | 0.87 | 1564587 | 0.95 | 1564587 | 0.95 |

**Table 8.** Genotype Prediction Report.

|  | *Precision* | *Recall* | *f1-score* | *Support* |
| --- | --- | --- | --- | --- |
| **Homozygous** | 0.75 | 0.82 | 0.78 | 11 |
| **Hetrozygous** | 0.91 | 0.75 | 0.82 | 28 |
| **Non-Variant** | 1.00 | 1.00 | 1.00 | 3954 |
| **Non-SNP** | 0.8 | 0.57 | 0.67 | 7 |
| **avg/total** | 1.00 | 1.00 | 1.00 | 4000 |

**Fig. 16.**
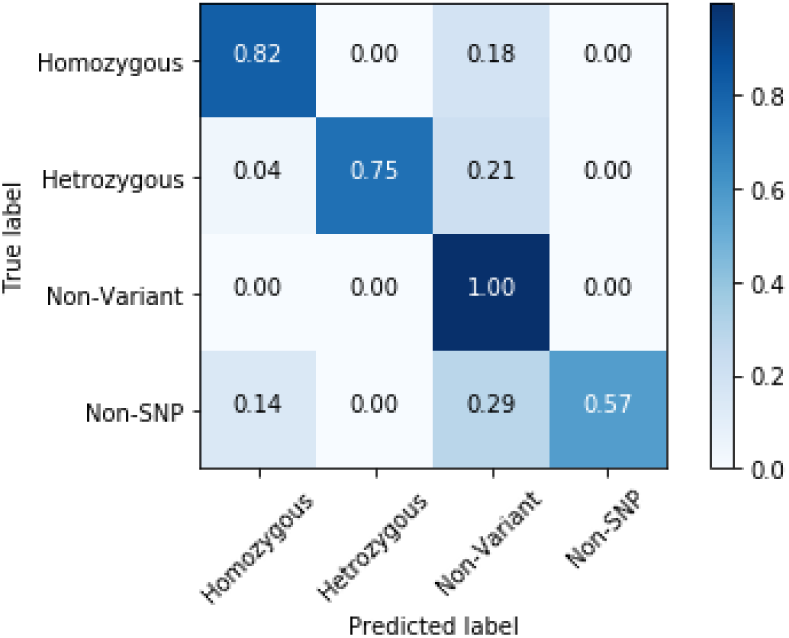
Confusion Matrix of Genotype Prediction.

#### 4.2.3 Experimental Setup and Results

CNN is modeled using *Tensorflow* library [29] on Ubuntu 16.04 operating system, which helps in scaling CNN network on both CPU and GPU based architecture. For benchmarking GATK performance *GATK 3.7* with *JAVA 8* is used. The hardware specification of system used for benchmarking is as shown in Table 9 :-

**Table 9.** System Specification.

| **Processor** | **Memory** | **Hard Disk** | **GPU** |
| --- | --- | --- | --- |
| Intel® i7-4770 CPU @ 3.40GHz | 7.7 GB | 1.5 TB | None |

We benchmark the performance of our variant caller network considering real execution time (in seconds) and accuracy of prediction. We have carried out comparative study of the proposed model against GATK 3.7 Haplotype Caller for “Illumina” dataset. To prove that deep learning based variant caller can be used across different NGS technologies we have also done performance and sensitivity analysis on “PacBio RSII” dataset.

We have used Dataset as mentioned in Table 10.

**Table 10.** Variant Caller Dataset.

| **Dataset** | **Species** | **NGS** | **Reference** |
| --- | --- | --- | --- |
| hg38.NA12878<br>chr21-14069662-46411975 | Homo sapiens | PacBio<br>RSII | hg38.chr21 |
| hg38.NA12878<br>chr22-18924717-49973797 | Homo sapiens | PacBio<br>RSII | hg38.chr22 |
| NA12878<br>chr20.10_10p1mb | Homo sapiens | Illumina | ucsc.hg19<br>chr20 |
| SRR1610046_1 | Ecoli | Illumina | ecoli.fa |
| SRR1610046_2 | Ecoli | Illumina | ecoli.fa |
| SRR1610047_1 | Ecoli | Illumina | ecoli.fa |
| SRR1610052_1 | Ecoli | Illumina | ecoli.fa |

**Table 11.** CNN Variant Caller vs GATK Haplotype Caller.

| Dataset | Number of Reads | Variant Caller Preprocessing Time(s) | Variant Caller Prediction Time(s) | Variant Caller Total Time | GATK Haplotype Caller Time(s) |
| --- | --- | --- | --- | --- | --- |
| NA12878-chr20 | 52053 | 12.17 | 0.75 | 12.92 | 20.2 |
| SRR2724094_1 | 513785 | 21.67 | 4.05 | 26.576 | 1745 |
| SRR2724094_2 | 341788 | 21.67 | 4.139 | 25.809 | 1716 |
| SRR2724099_1 | 1535766 | 90.52 | 16.89 | 107.41 | 2830 |
| SRR2724099_2 | 1527685 | 65.611 | 10.64 | 76.251 | 3322 |

For Human data we have used four different datasets for benchmarking, *GRCh3718* data is used as reference genome. Network is trained using first 30000 images of *NA12878-chr21* dataset and validated with next 40000 images. Trained network is used for variant and indel prediction of *NA12878-chr22* and *NA12878-chr20* datasets. For non-human data, we have used EColi data. Model is trained using *SRR1610046_1* EColi dataset, and tested with four other Ecoli datasets (Table 10). Timing analysis for *NA12878-chr21, NA12878-chr22* and *SRR1610046_1* datasets is shown in Fig. 17. Prediction accuracy of CNN Variant Caller when trained and tested on same species varies between 86%-93% compared to true variant as shown in Fig. 18. To observe accuracy across species, we have used Variant Caller trained on Human Genome *NA12878-chr21* and have used it for predicting variant in ecoli dataset *SRR1610046_1*. We have compared the prediction accuracy when trained on dataset from same species to dataset of different species. Results are comparable as shown in Fig. 19.

**Fig. 17.**
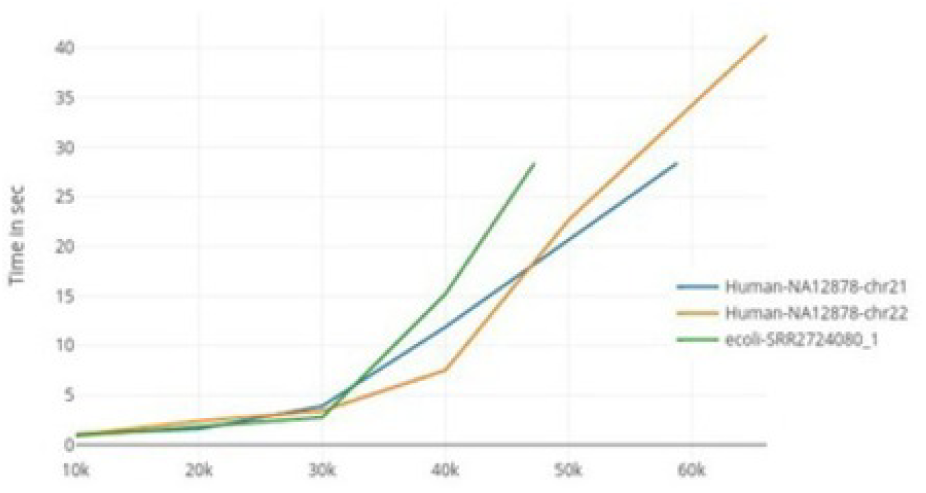
Variant Caller Timing Analysis.

**Fig. 18.**
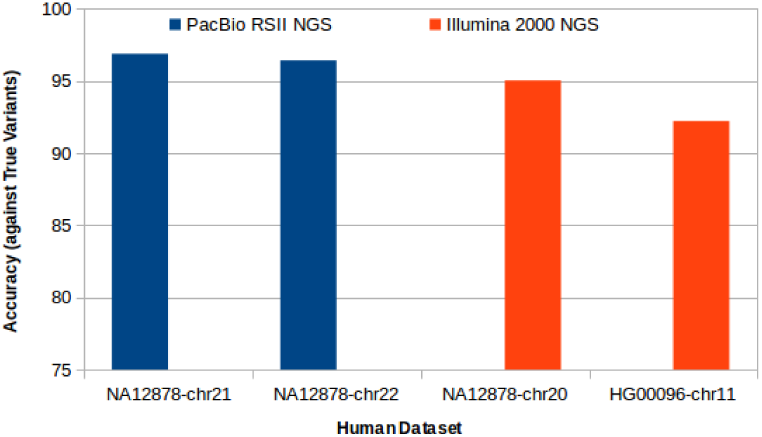
Prediction accuracy of Variant Caller Across NGS Technologies.

**Fig. 19.**
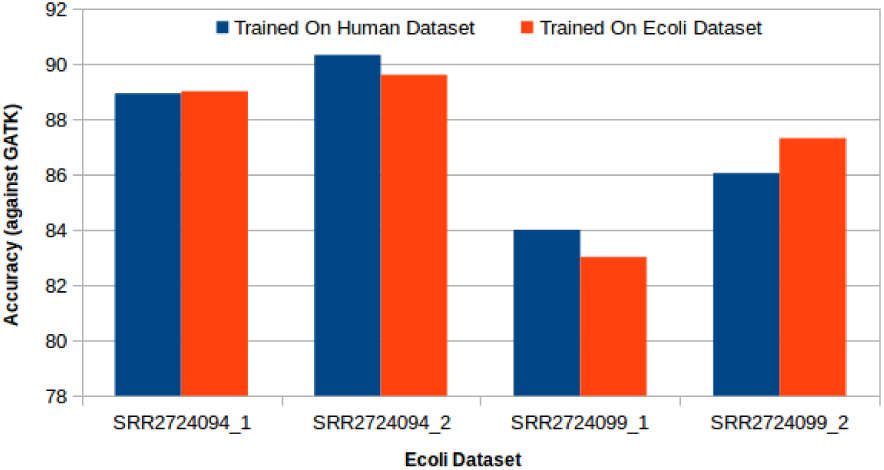
Prediction Accuracy if trained with same species dataset vs different species dataset.

Moreover, we tried to analyze how the model trained on one NGS technology can be used across other tecnology. In this test we have used variant caller trained on human *NA12878-chr21* of PacBio RSII data and predicted accuracy on PacBio RSII dataset of human *NA12878-chr21* and *NA12878-chr22* and on Ilumina 2000NGS human dataset *NA12878-chr20* and *HG00096-chr11*. The variant prediction accuracy in both test cases is above 90 % as shown in Fig 18. Hence variant caller trained on dataset of any species can be used for variant prediction of any other species. This will allow non-human genome sequencing project to be benefited by depth of human truth data[**?**]. And Model Trained on one NGS technology can be used across various other NGS. Thus moving away from expert driven model to more generic data driven model.

## 5 CONCLUSION AND FUTURE WORK

In this paper we have explored various deep neural network based models to tackle two problems in Genome Sequencing-namely, Global/Local Alignment of raw NGS data and Single Nucleotide Variant(SNV) Identification and its Classification. Proposed solutions were tested and their timing performance were benchmarked against existing de-facto standard tool sets. We have observed that performance of our alignment model is comparable to BLAST but is slower than BWA-mem. Our DNN based SNV model at preliminary level was found to perform faster than GATK 3.7 and is highly specific. There can be further boost in performance of these DNN models by using more biological information and by using multi-process and multi-threaded programing models. We have also established that use of DNN makes our model independent of underlying NGS technologies and model trained for single NGS technologies can be used across all others. Thus with help of Deep Neural Network we can generate models which provide faster,reliable and highly specific solution for identification of SNV from raw NGS reads, and will pave path for data centric solutions as compared to expert based methods in field of bio-informatics.

## A ALGORITHMS

### Algorithm 2 Data Generation

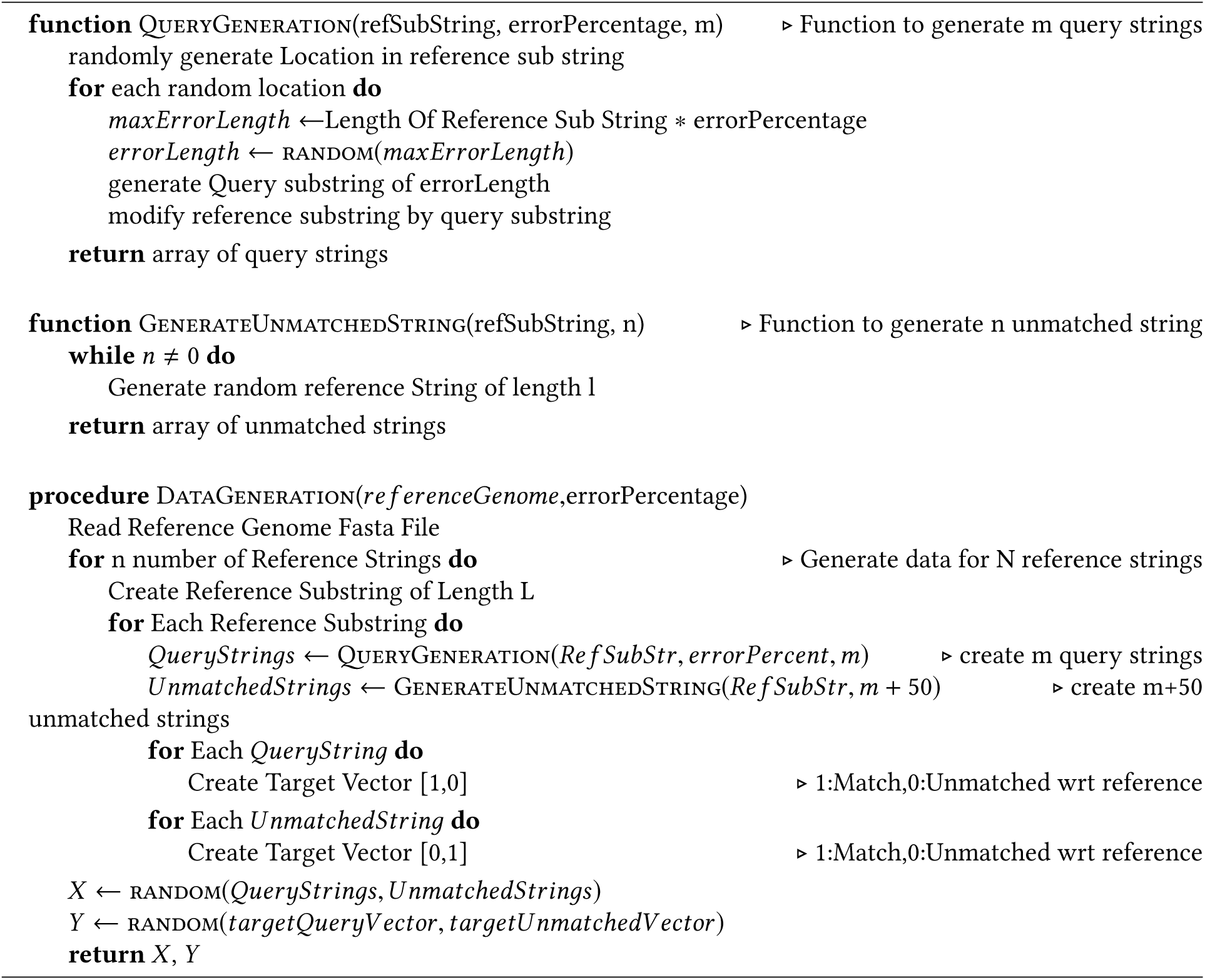

### Algorithm 3 Global Alignment Using Deep Conflation Model

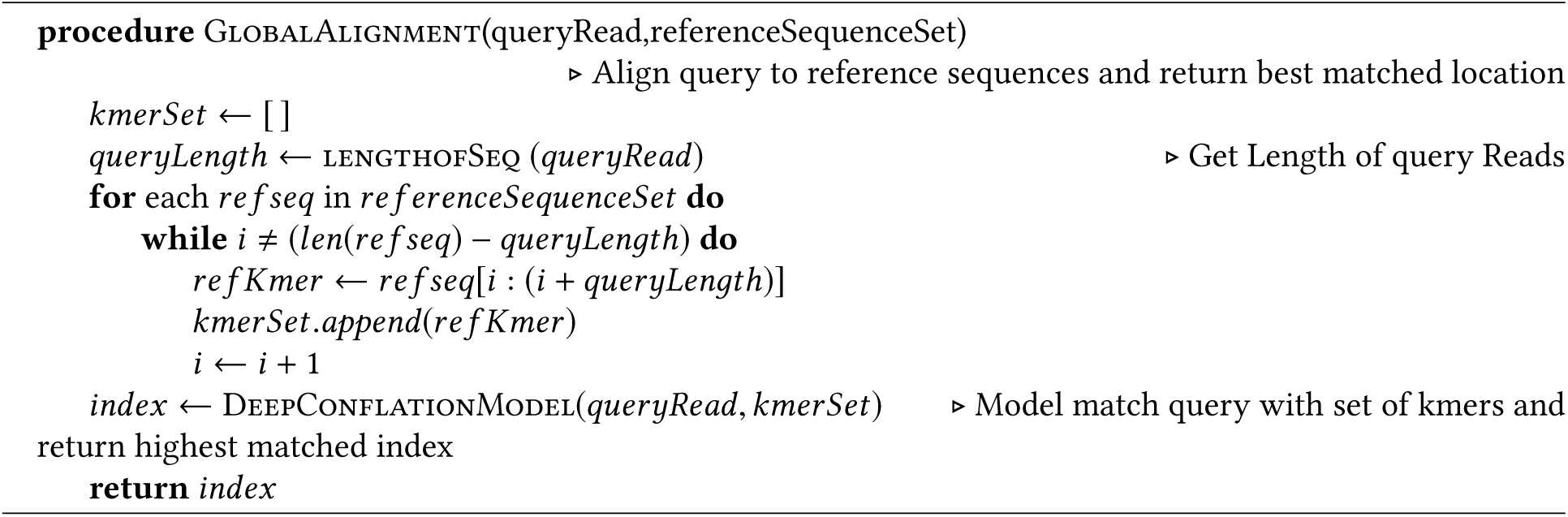

### Algorithm 4 Population Evolution

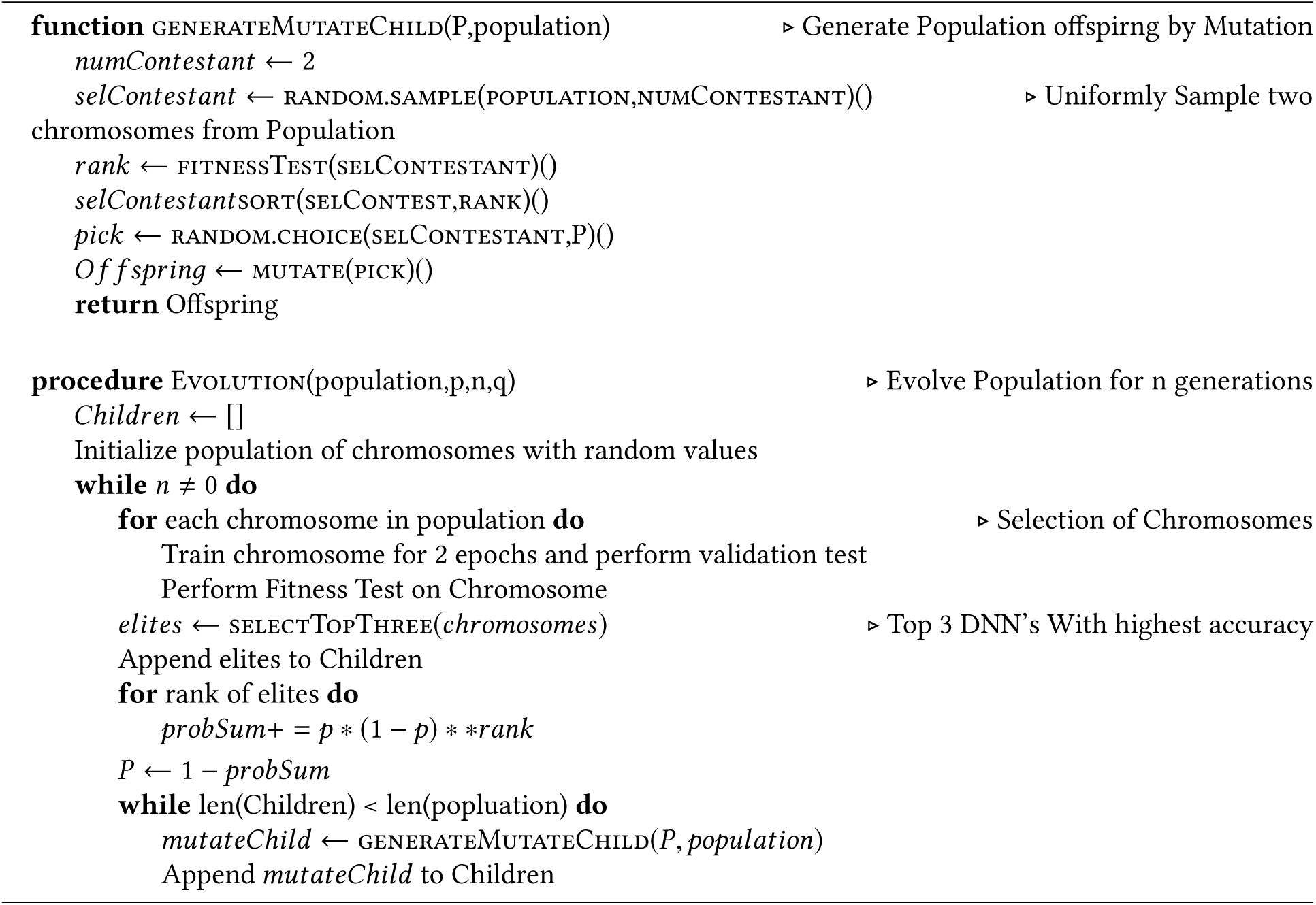

### Algorithm 5 Local Alignment Using RNN Model

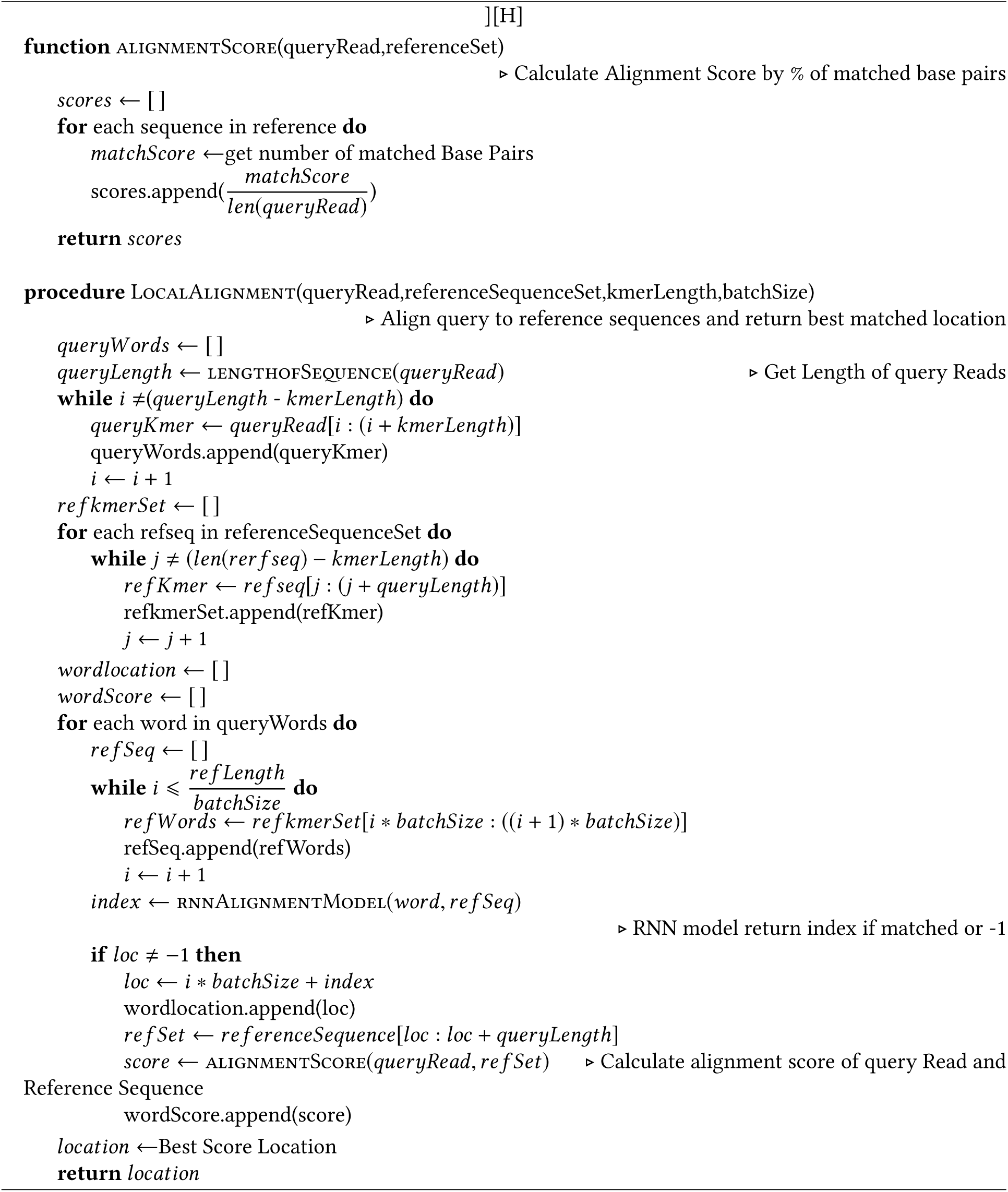

## B EXPERIMENT DETAILS OF DEEP CONFLATION MODEL

### B.1 Simulated Data Generation

Simulated data is generated by adding random errors in the form of substitution and indels of bases of corresponding reference string. Through this scheme, we intend to generate data which has close resemblance with actual reads obtained from NGS sequencer. Also, by using error as a parameter in the data generator, we can generate and experiment with datasets containing varying degree of errors. This will help in establishing correlation between the percentage of error in reads and the prediction accuracy of Neural Network, if any exists. Algorithm for data generation is described in Algorithm 2 (Appendix A).

The output of data Generation is jumbled using uniform random distribution. For experimentation, three data sets of inputs and targets are generated - namely, training dataset, validation dataset and test dataset. Each dataset is generated by method as described above, but the number of matched and unmatched corpuses are different due to uniform random sampling of batch data. Data is generated for 4000 reference strings, total database of corpuses is of the order 61* 4000. Dataset is split into 80, 20 ratio for training and validation set. For test, another 1000 reference string dataset is generated. Length of each reference string is varied from 15-170 bps for experimentation. Details of DNN models which use this dataset are discussed in following section.

### B.2 Deep Conflation Model for Alignment

#### Model Overview

Given a read k-mer ‘x’, the model ranks a set of reference k-mers < *r*_1_, *r*_2_,, *r*_*n*_ >, so that the reference k-mer most similar to given read gets highest rank. The k-mers are first preprocessed and encoded as per the following scheme:

A : 0, C : 1, G : 3, T : 4.

The model consists of two parts:

1. Extracting finite dimension feature vectors from the encoded read and reference k-mers through CNN.
2. Ranking the reference features based on cosine similarity with read features.< *r*_1_, *r*_2_,, *r*_*n*_ > are ranked in order of decreasing similarity.

#### CNN Feature extractor

Let ‘x’ be a string containing ‘k’ characters. Then, character encoding of x is obtained as *x* = [*x*_1_, *x*_2_…, *x*_*k*_] x is then fed as input to the CNN, and convolution is applied using three different filters of sizes 2, 3 and 4 to extract features corresponding to bigrams, trigrams and 4-grams. The ‘t’th output of a convolution with filter size ‘f’ is obtained as follows:

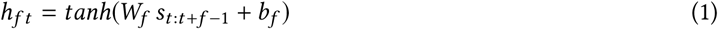

Where *W*_*f*_ is the convolution weight, *b*_*f*_ is the bias and *s*_*t* :*t* +*f* − 1_ is the vector obtained by concatenating *s*_*t*_ to *s*_*t* +*f* − 1_ Feature map of convolution with filter size ‘f’ can be defined as :

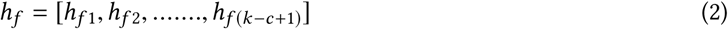

Max pooling is applied to feature map *h*_*f*_ to obtain *ĥ*_*f*_. Feature maps obtained with filters of sizes 2, 3 and 4 are concatenated to form a vector representing the entire input string/character sequence *y*= [*ĥ* _2_, *ĥ* _3_, *ĥ* _4_] This final feature vector is fed to the ranker module. The process of feature extraction is repeated for each read k-mer and reference k-mer.

#### Ranker

After obtaining the feature vector *y*_*x*_ for read k-mer x, and set of feature vectors; 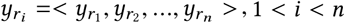 for reference k-mers < *r*_1_, *r*_2_,, *r*_*n*_ > the ranker computes cosine similarity between *y*_*x*_ and each of 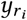 as follows

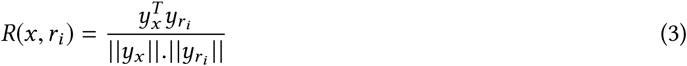

Reference k-mers are ranked by these scores with respect to a given read k-mer, with the best matching reference k-mer awarded the highest rank.

#### Training

For training, we have used *simulated data* as described in previous subsection. We feed the model a batch of 100 such *(reference,mutant)* pairs, wherein the model is expected to learn to match the mutated string to its original reference string, and assign it the highest rank. Using the similarity scores obtained from equation 3, the posterior probability of correct reference string, given the mutant string is defined as follows:

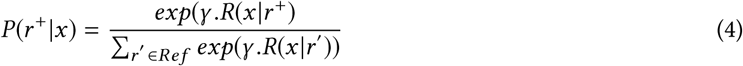

Where *r* ^+^ is the correct reference string to which x should match. *γ* is a hyperparameter set to 10. *Ref* is the set of reference strings, containing *r* ^+^ and 99 other random non matching reference strings. While learning, the model tries to minimize the following loss function:

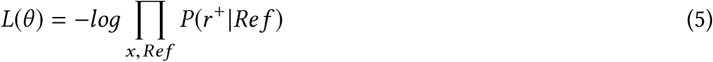

Where *θ* denotes the model parameters to be learnt, and product is over all examples in a batch. Adam Optimizer with learning rate of 0.0002 is used for loss minimization.

#### Testing

During testing, a read k-mer is fed to the model and features extracted from this input are compared with feature vectors obtained from reference k-mers. Thereafter, reference k-mers are ranked in order of similarity (based on Equation 3), and the highest rank reference k-mer is output as the most probable match for the given read k-mer.

## C DESIGN AND TRAINING OF RNN MODELS

### C.1 Vanilla RNN

A vanilla RNN is designed to include an additional context layer, where the activation of hidden units from previous feed-forward process is stored [30]. These hidden state values are concatenated with input which generates three dimensional vector of shape [*sequence*_*length, batch*_*size, hidden*_*size* + *input* _*size*] These vectors are then fed to RNN cell with fully connected nodes. Each cell of RNN stores value as hidden state which is of dimension [*batch*_*size, hidden*_*size*] The output of RNN cell is then fed to FC layer while hidden state is fed to input layer of next cell.

#### C.1.1 Training

The network was trained on *simulated data* for 50 epochs, with 380 batches of size 10 (Number of batches = 61* 4000/batch size/Sequence length) for each epoch. For each epoch, at interval of 100 batches, training loss and training accuracy of predicted output was calculated. To observe overfitting, validation test was performed on validation data at same interval. With learning rate of 0.001 for optimizer the plots of loss function, training accuracy and validation accuracy is shown in Fig. 20

**Fig. 20.**
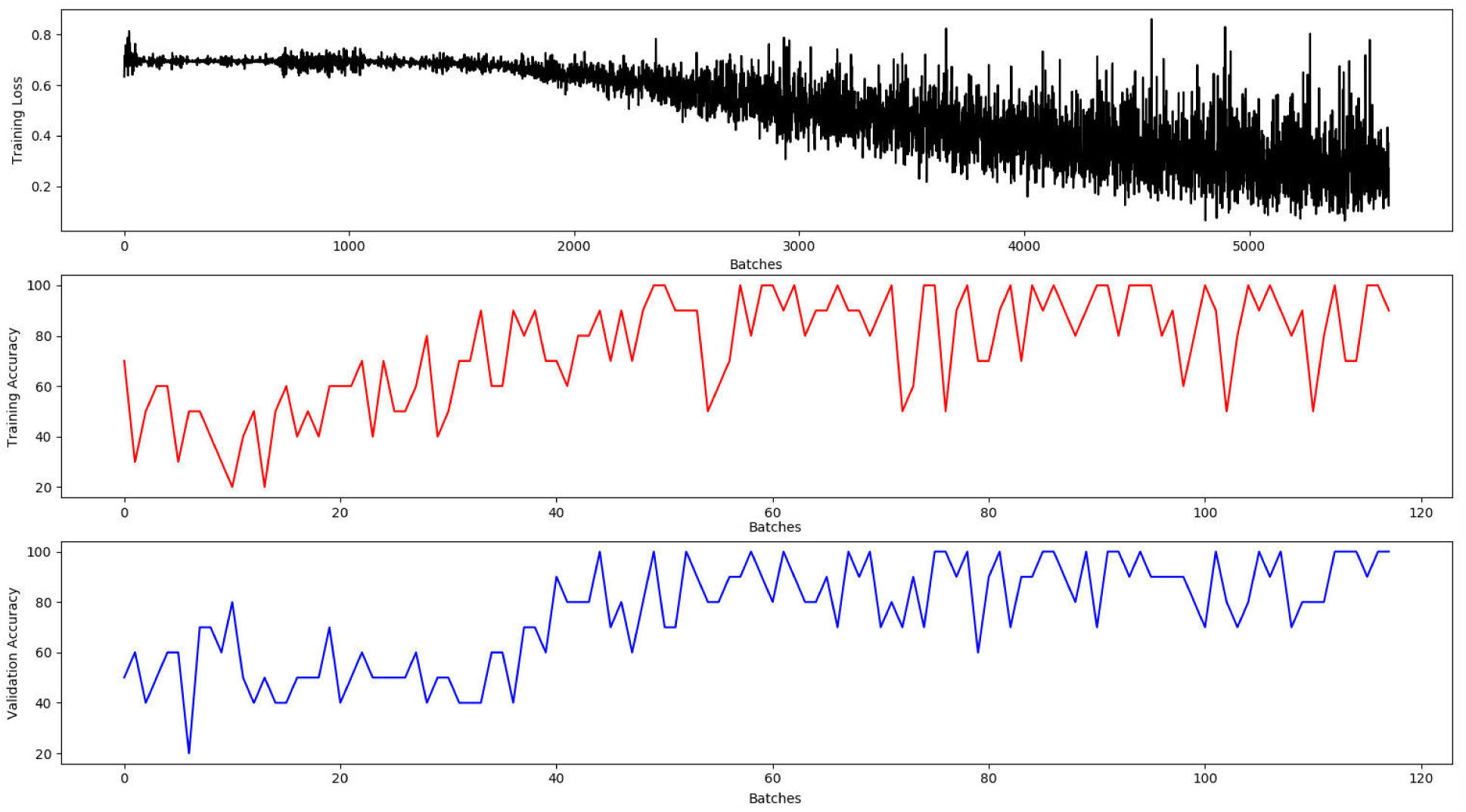
Training Loss, Training and Validation Accuracy Of Vanilla RNN.

### C.2 LSTM

In order to model global alignment of genome sequences by comparing reads of long lengths, we have explored LSTM cell in our Recurrent Neural Network architecture. Benefit of LSTM is that it helps to avoid long-term dependencies and thus remembers data patterns for long time(sequences). From many different LSTM configurations available, we have used Hochreiter & Schmidhuber LSTM [31]. A cell of LSTM consists of hidden state and cell state. *Hidden state* of LSTM cell is used for modifying state of next LSTM cell, while *Cell State* dictates output of current cell. Both values for initial LSTM cell are initialized to zero. Similar to RNN, input to cell is concatenated with previous hidden state before being fed to next LSTM cell.

#### C.2.1 Training

LSTM network was trained for 100 epochs with number of batches and batch size same as that of RNN model. Training set and validation set are also kept same as that of RNN model. Unlike RNN, the training loss and training accuracy in LSTM model are calculated for each epoch, while validation test is done at an interval of 100 batches for each epoch. The training loss, training accuracy and validation accuracy of LSTM model is shown in Figure 21

**Fig. 21.**
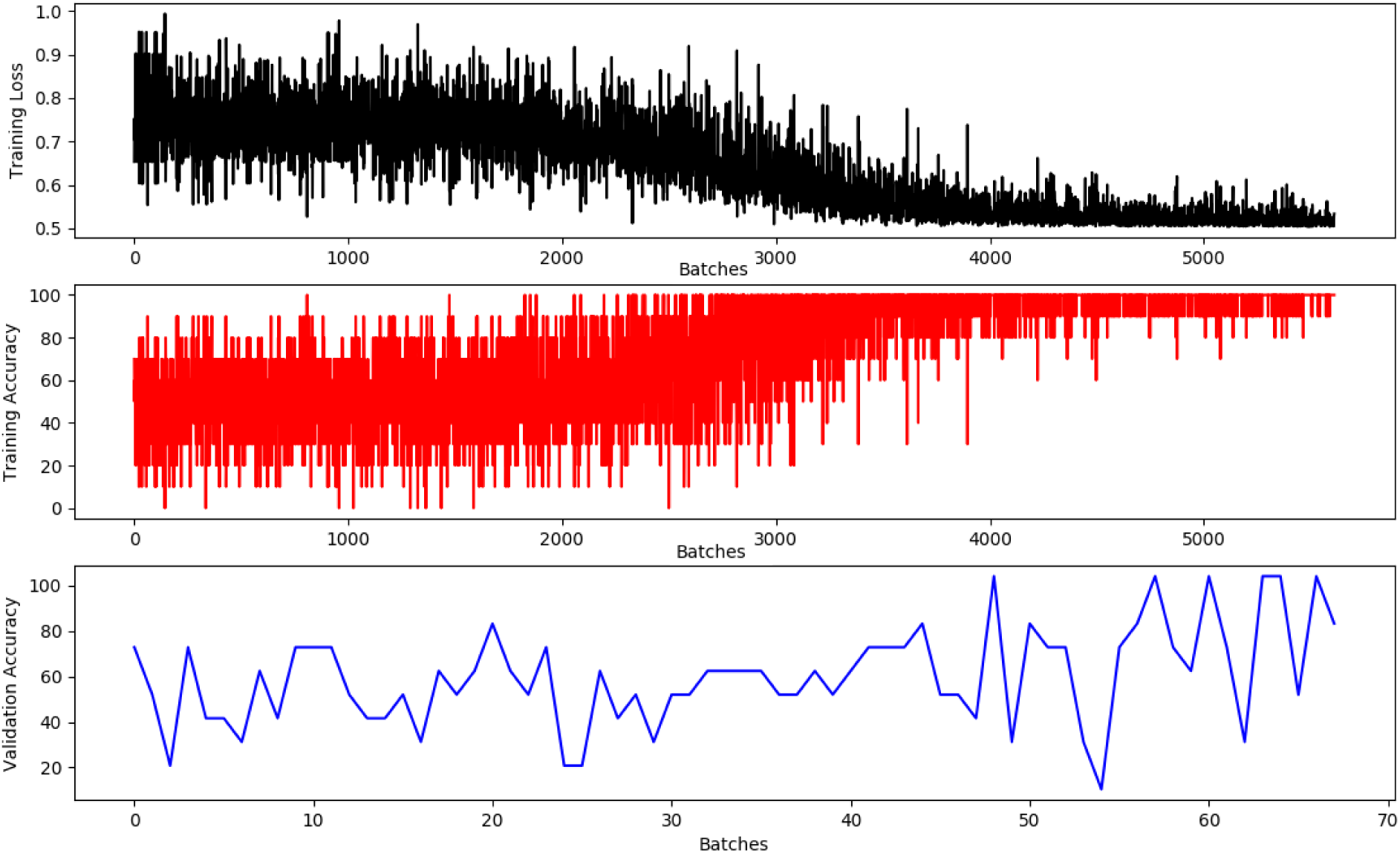
Training Loss, Training and Validation Accuracy Of LSTM.

## D DATA-PREPROCESSING FOR CNN BASED VARIANT CALLER

### D.0.1 Data Preprocessing

This process involves generation of variant candidates and generation of labeled data to train CNN. For variant generation, the aligned BAM file is read using Samtools and corresponding CIGAR [32] strings are decoded. From CIGAR string, different alleles along with their position in reference reads are identified by comparing it with reference read. These alleles are then classified either as reference-matching base, reference-mismatching base, an insertion or as a deletion [13].

The frequency of each distinct allele for a position is calculated using Position Specific Frequency Matrices(PSFM) [33] as shown in Fig.22. If frequency is above set threshold, the allele site is considered as a variant candidate. For all possible variant candidates, Variant string is generated by reading fix length of bases from reads, with variant site as median. These variant candidate strings are encoded into three matrices. 1^st^ matrix is created by encoding the expected reference sequence using one-hot-like encoding. It encodes number of reads that aligned to a reference position. The 2^nd^ matrix encodes the difference of all the bases observed in the read-reference alignment. Reads with ‘N’ nucleotides are ignored. The 3^rd^ matrix is similar to the 2^nd^ matrix, except, none of the insertion bases in the reads is counted. All the matrices are input to CNN for both training and variant calling. Fig.23, depicts data transformation at each stage of data preprocessing.

**Fig. 22.**
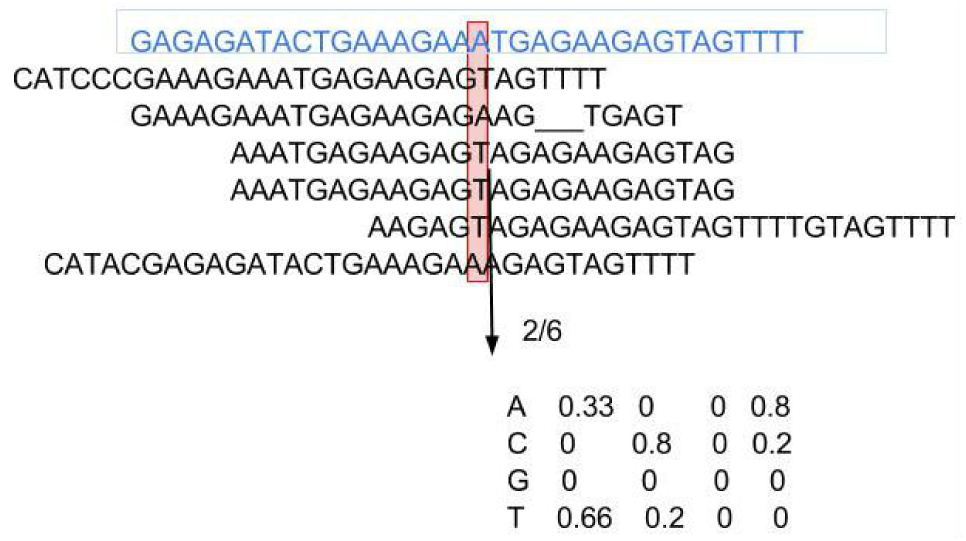
Position Specific Frequency Matrix.

**Fig. 23.**
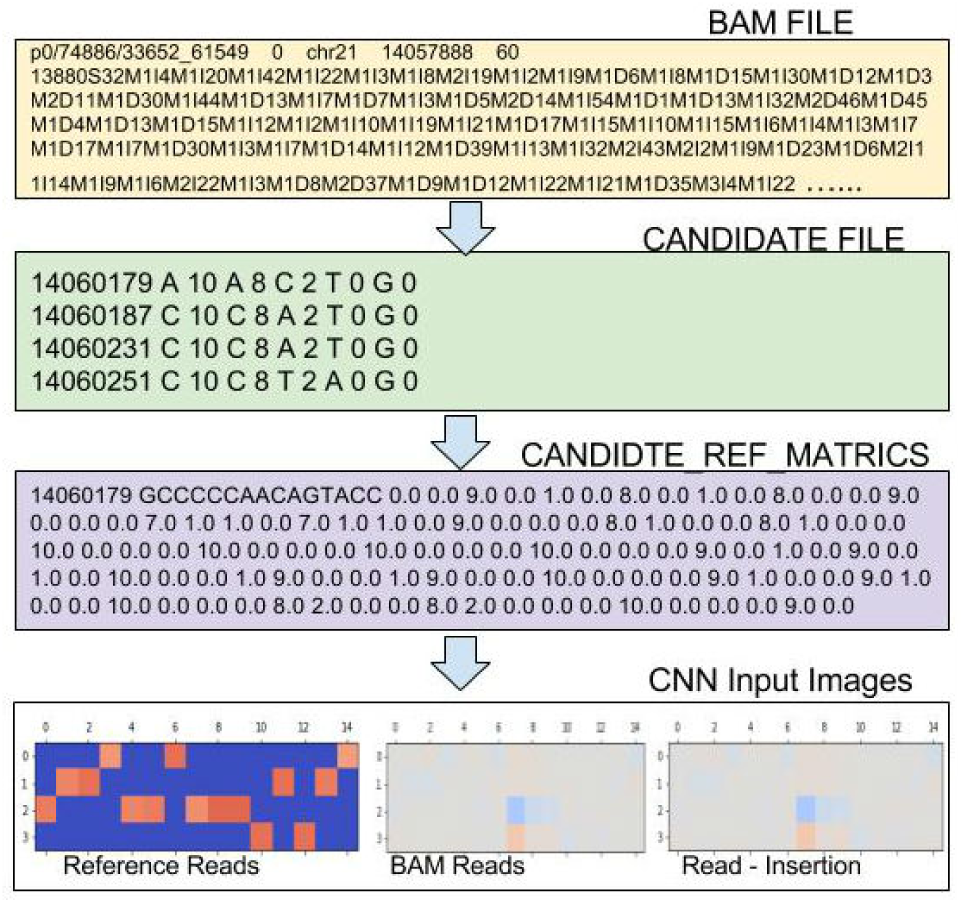
Data transformation during Preprocessing.

#### Labeled Data

Labeled Data is created by reading corresponding variant file in,VCF format. Only those true variants are considered for labeling which fall in high confidence region of reference genome. For all true variants, their position,reference base, alternate base and their genotypes are read. These variants and their genotypes are then encoded as shown in Fig. 24b.

The process of generating label data and data transformation for labeling is shown in Fig.24a.

**Fig. 24.**
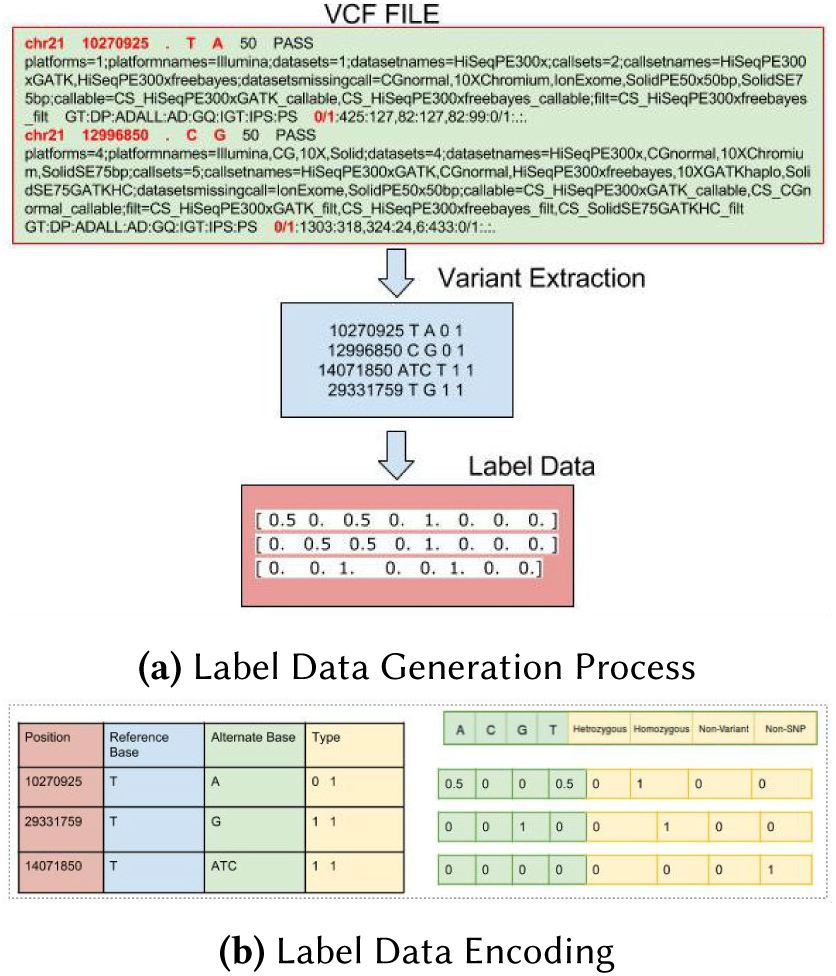
Label Data Generation.

